# Structure of *Dunaliella* Photosystem II reveals conformational flexibility of stacked and unstacked supercomplexes

**DOI:** 10.1101/2021.11.29.470333

**Authors:** Ido Caspy, Maria Fadeeva, Yuval Mazor, Nathan Nelson

## Abstract

Photosystem II (PSII) generates an oxidant whose redox potential is high enough to enable water oxidation^1,2^, a substrate so abundant that it assures a practically unlimited electron source for life on earth^3^. Our knowledge on the mechanism of water photooxidation was greatly advanced by high-resolution structures of prokaryotic PSII^4–6^. Here we show high-resolution structures of eukaryotic PSII from the green algae *Dunaliella salina* at two distinct conformations. The conformers are also present in stacked PSII, exhibiting flexibility that is relevant to the grana formation in chloroplasts of the green lineage. CP29, one of PSII associated light harvesting antennae, plays a major role in distinguishing the two conformations of the supercomplex. We also show that the stacked PSII dimer, a form suggested to support the organization of thylakoid membranes^7,8^, can appear in many different orientations providing a flexible stacking mechanism for the arrangement of grana stacks in thylakoids. Our findings provide a structural basis for the heterogenous nature of the eukaryotic PSII on multiple levels.

## Introduction

In eukaryotes, the light reaction of oxygenic photosynthesis occurs in chloroplasts. Four protein complexes essential for the light reactions reside in an elaborate membrane system of flattened sacs called thylakoids^9^. From these four complexes the Photosystem II (PSII) complex catalyzes light-driven water oxidation and provides the electrons used for carbon fixation^10,11^.

Thylakoids form a physically continuous three-dimensional network, differentiated into two distinct physical domains: cylindrical stacked structures (called grana) and connecting single membrane regions (stroma lamellae). Photosystem I (PSI) is mainly located in the stroma lamellae while PSII is found almost exclusively in the grana^12–14^. Grana stacking is a dynamic process dependent on the internal osmotic pressure, the luminal ion composition, and environmental ques, and is thought to be supported by interactions among PSII complexes^15–19^.

PSII is a homodimer with a molecular mass of ∼500 kDa, each monomer contains cofactors such as chlorophylls (Chls), quinones, carotenoids, and lipids which are coordinated by at least 20 protein subunits^1,20^. In each PSII core, a cluster of four manganese (Mn) and one calcium (Ca) carries out H_2_O oxidation and O_2_ release^21^. The eukaryotic reaction centre (RC) is a dimer surrounded by tightly bound monomeric light harvesting complexes (LHCs) and trimeric LHCII complexes^22–24^; Two monomeric LHCs, CP26 and CP29, are located between LHCII trimers and PSII core subunits^25^; additional LHCII trimers can bind PSII depending on light intensity and quality^26^.

Although more than three billion years of evolution separate cyanobacteria, red algae, green algae and plants, high resolution PSII structures show that each PSII monomer along with its dimeric arrangement are highly conserved, especially in the membrane bound regions of the PSII^27–32^. Structural and spectroscopic investigations uncovered various aspects of PSII’s water splitting mechanism, but a complete model is still missing^4,5,33–35^. Most of the mechanistic and structural studies of PSII were performed in thermophilic cyanobacteria, but structural studies of PSII from the eukaryotic lineage are lagging^27,31,32,36^.

In this work photosystem II was isolated from the halotolerant green alga *Dunaliella salina*. A high-resolution (2.43 Å) structure of PSII shows unique structural properties of the *Dunaliella* PSII supercomplex. The eukaryotic PSII appear to exist in two distinct core conformations that differ substantially in their inner dimer separation and the location of CP29, an important monomeric LHC. Structural analysis of stacked PSII dimers showed highly flexible interactions which can play a role in the dynamic organisation of chloroplast membranes. These finding introduce an additional, underlying, level of organisation which can impact its excitation energy transfer properties and the overall organization of the thylakoid membranes.

## Results

### Two distinct PSII conformation in green alga

Highly active PSII from *D. salina* cells was applied on glow-discharged holey carbon grids that were vitrified for cryo-EM structural determination (see Methods). Initial classification of the dataset showed that approximately 20% of the particle population were in a stacked PSII configuration, containing two PSII dimers facing each other on their stromal side (**Extended Data Fig. 1)**. From the unstacked PSII dimers, approximately 20% were in the C2S configuration (two Cores, one Stable LHCII), which was previously identified by low resolution cryo-EM^25^ (**Extended Data Fig. 1c**). The majority of the PSII particles contained 2 LHCII in the C2S2 configuration. The map of the C2S2 particles refined to a global resolution of 2.82 Å (**Extended Data Fig. 2)**. Close examination of this map revealed that CP29, one of the monomeric LHC proteins, implicated as a junction for excitation energy transfer from LHCII trimers to PSII core, appeared to be in lower resolution than the rest of the supercomplex (**Extended Data Fig. 2a**). Indeed, when this particle set was further classified, two distinct PSII conformations of the PSII supercomplex became apparent. In these two conformations the two PSII cores are shifted latterly with respect to each other (**Fig. 1**). This lateral shift is accompanied by several other associated movements, most noticeably, a large movement of the CP29 subunit (**Fig. 1**) in line with our initial observation. The two conformers were denoted compact and stretched PSII (C2S2_COMP_ and C2S2_STR_, respectively), and the high specific activity of 816 μmol O_2_/mg Chl/h measured for the preparation prior to vitrification suggest that both are highly active. The final reconstruction of the compact orientation refined to an overall resolution of 2.43 Å, the highest of any eukaryotic PSII structures (PDB ID 7PI0; **Fig. 1a** and **Extended Data Fig. 1-3**).

**Fig. 1:**
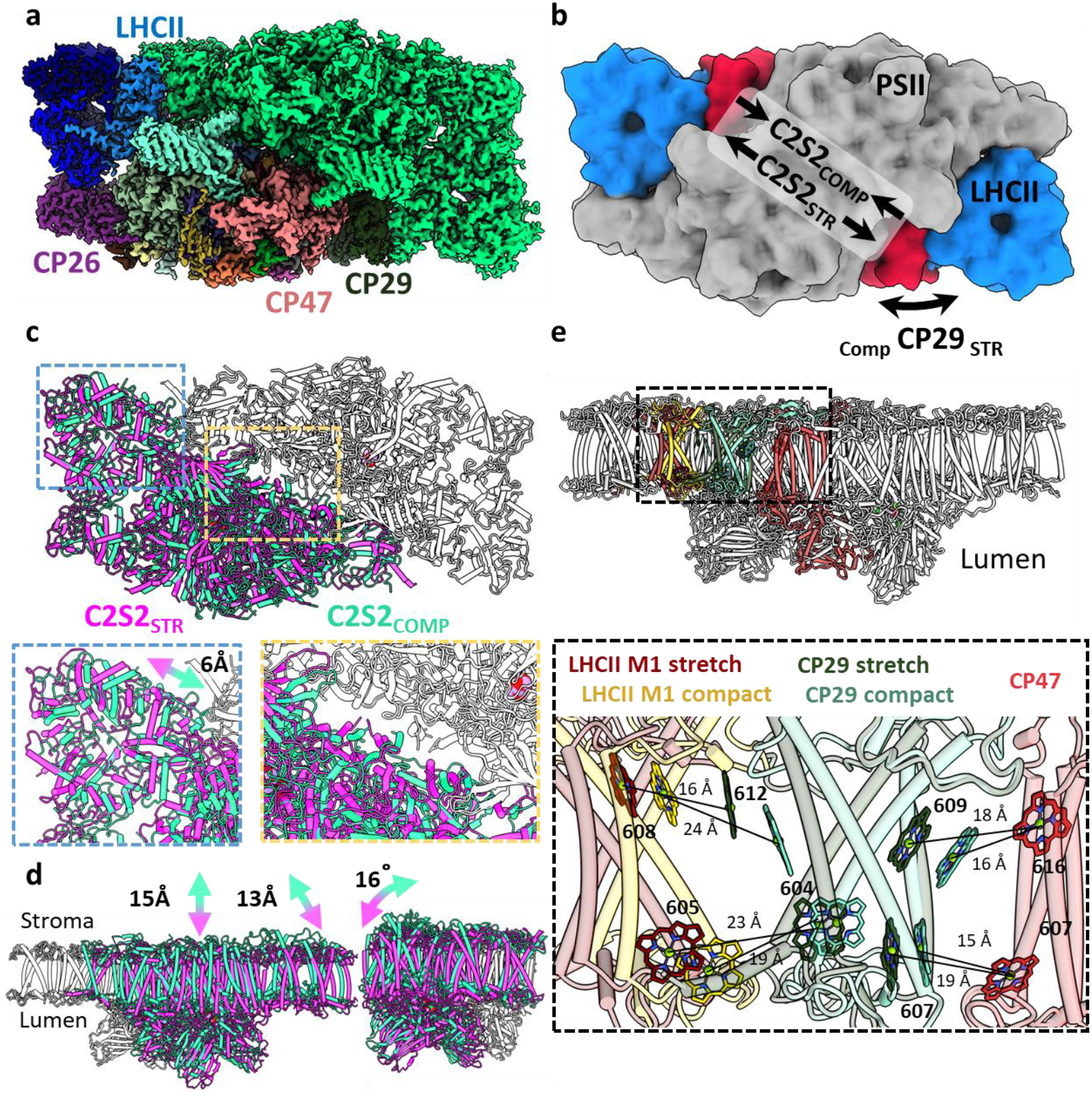
Two conformation of the eukaryotic PSII. **a**. overall view of the PSII C2S2 map in the compact conformation. One asymmetric unit is colored in green, in the other each chain is colored individually. **b**. Low resolution model depicting the overall shifts in subunits between the two PSII conformations. CP29 in red, LHCII in blue and the two PSII cores in grey. **c**. The two PSII conformations were superposed on one asymmetric unit (colored in grey). The second asymmetric unit is colored in magenta for the stretched conformation (C2S2_STR_) and green for the compact conformation (C2S2_COMP_). A close up showing a 6 Å shift in the position of LHCII and the lateral displacement between the two cores. **d**. The stretched PSII conformation shows substantial drop in the membrane plane (13 to 15 Å, depending on the precise location), contributing to a larger inward curve (compared to the luminal space) of the entire supercomplex. Large deformations in the position of CP26 subunit which rotates by 16° between the two conformations. **e**. Considerable changes in the position of CP29 affect the transfer rates between LHCII and CP47. CP47 of both conformations (in red) is superposed and distances between key Chls of CP29_COMP_ (light turquoise) and CP29_STR_ (dark green) shows increased transfer distances in the stretched conformation. The distances between LHCII and CP29 follow an opposite trend, decreasing in the stretched conformation (LHCII_STR_ in dark red) and increasing in the compact conformation (LHCII_COMP_ in yellow).

The C2S2_COMP_ structure is similar to the previously determined C2S2 supercomplex from *Chlamydomonas reinhardtii* or higher plants^31,36^ and the cyanobacterial core structures^4,6^. The second, stretched conformer, was solved to 2.62 Å resolution, and accounted for about 37% of the unstacked C2S2 PSII particles. **Fig. 1** and **Extended Data Movies 1-2** depict the superposition of the polypeptide chains of the two conformers, showing major differences in the location and orientation of PSII monomers. Superposition of the *Dunaliella* and *Chlamydomonas* C2S2 (PDB 6KAC) structures and maps suggests that the *Chlamydomonas* structure also contains these different conformers. This may explain the decreased local map resolution presented in the aforementioned subunits, compared to the rest of the cryo-EM map^31^.

### Structures of *Dunaliella salina* unstacked PSII at high-resolution

Thus far, available PSII structures suggested a single, highly conserved organisation of the two PSII cores^4,27–29,31,32^. The high-resolution structures of *Dunaliella* C2S2_COMP_ and C2S2_STR_ provide a new perspective on the dynamic arrangement of eukaryotic PSII and the interaction of the core complex with its LHCs. To compare the C2S2_COMP_ and C2S2_STR_, the core complexes were aligned (**Fig. 1 and Extended Data Movies 1-2)**. Initial inspection showed that one of the major differences between the two conformations is the orientation of CP29 (**Fig. 1e**). In C2S2_STR_ CP29 helices I and III move towards the LHCII trimer of the opposite monomer and away from CP47, with helix II of CP29 serving as a rotation axis. Moreover, CP29_COMP_ contained only 9 Chls, compared to 11 in CP29_STR_ and13 Chl in *Chlamydomonas* PSII CP29^31^. Chls 605 and 616 were absent in both structures and CP29_COMP_ was also missing Chls 611 and 613. This might be attributed to the flexibility of CP29 C-terminus, or the internal rearrangement caused by its movement.

In the C2S2_STR_ conformation the PSII monomers slid in the membrane plane along the central symmetry axis separating them (**Fig. 1b** and **Extended Data Movies 1-2**). The non-aligned core shows the extent of the shift in the core peptides together with the minor LHCs and LHCII trimer (**Fig.1c** and **d**). As a result, all the interactions at the cores interface are modified, leading to local changes in chain orientations and the conformations of some loops. Core subunits at the centre of the monomer displayed a greater shift (D1, D2, CP47 CP43 and PsbO were displaced by 6-10 Å; **Fig. 1**), and the peripheral subunits showed the largest shift and tilt compared to C2S2_COMP_ (PsbE, PsbP and CP26 moved by 13 Å, PsbZ showed the largest relocation of nearly 15 Å. and CP26 showed a maximal tilt of 16°; **Fig. 1**). Multibody refinement^37^ of both C2S2_COMP_ and C2S2_STR_ demonstrated that the two PSII monomers in each conformation contain additional structural heterogeneity (**Extended Data Fig.4-5** and **Extended Data Movie 3-6**).

### Distinct CP29 conformations alter LHCII to PSII core connectivity

The observed conformational change of CP29 alters excitation energy transfer (EET) pathways from LHCII to the PSII core and may account for the differences between calculated and measured EET^23,38–41^. To assess changes in transfer rates between the stretched and compact orientations, we measured how the distances between the closest Chls of CP47 (PSII core), CP29 and LHCII change between the two PSII conformations. The average distances between CP29 Chls 603, 607 and 609 to the CP47 Chls 607 and 616, changed from 17 Å to 20 Å between the compact to stretched conformations, suggesting that faster transfer rates from CP29 to CP47 in the compact conformation. In contrast to this, the distances between CP29 Chls 604 and 612 to LHCII Chls 604 and 608 increased from 20 Å in the compact conformation to 23 Å in the stretched conformation, suggesting that transfer from LHCII to CP29 is slower in the compact orientation. The missing CP29 Chls 611 and 613 form part of the interface to LHCII and are missing in the compact conformation, which should also contribute to slower transfer rates from LHCII to CP29 in the compact configuration (**Extended Data Fig. 6)**. Altogether, transfer from LHCII to the PSII core should be considerably slower in the compact orientation from both distance and Chl occupancy considerations. Similar features of altered Chl conformations were identified in molecular dynamics (MD) simulation of LHCII exploring its structural dynamics^42^ compared to its crystal structure. The analysis showed differences in the excitonic coupling of Chl clusters 606-607 and 611-612. MD suggested an increase in the interaction energies of 606-607 and a decrease in the interaction energies of similar proportion in 611-612^42^. The 611-612 Chl pair was proposed as a light-harvesting regulator of excitation energy transfer from CP29 to CP47 and as a quenching site^38^, as its change in fluorescence yield was attributed to a protein conformational change that leads to a redistribution of the interpigment energetics^43^.

### The compact and stretched PSII conformation contain substantial levels of continuous structural heterogeneity

Using multibody refinement^37^, with each PSII monomer defined as a separate rigid body, significantly improved the resolution and map quality in both C2S2_COMP_ and C2S2_STR_, showing that substantial structural heterogeneity exists in both datasets at the level of PSII monomers. Analyzing the shape of the heterogeneity in C2S2_COMP_ and C2S2_STR_, using Principal Component Analysis (PCA) showed that the first six Principal components (PCs) explains more than 85% of the variance in the data and consists of continuous heterogeneity (**Extended Data Fig. 4-5; Extended Data Movies 3-6**). Substantial displacements of approximately 13 Å are observed between the two monomers in the compact conformation (**Extended Data Fig. 4**) and a larger range of displacements (up to 20 Å) exists in the stretched conformation (**Extended Data Fig. 5**). The direction of PCs describes translations perpendicular and parallel to the membrane plane. This suggests that both conformations are flexible and can respond to different membrane curvature (**Extended Data Movies 3-6**). To examine the possible effects on energy transfer we measured the change in intermonomer Chl distances across the different components. As expected, the PCs describing changes in the membrane plane significantly change some key distances between LHCII, CP29 and D1 across monomers (**Fig. 2**). This means that within each PSII conformation, substantial levels of heterogeneity in transfer rates should be considered. Changes in Chl positions were observed in CP29 Chls linking CP29 to PSII core and those connecting CP29 with LHCII. These Chls moved by an average distance of more than 5 Å, in both conformations (**Extended Data Table 2**). This implies that the association between PSII monomers and between PSII cores and LHCs contains a certain degree of freedom which can modulate excitation energy transfer, the entire assembly may be affected by changes in thylakoid membrane properties such as fluidity, composition, and curvature^44,45^.

**Fig. 2:**
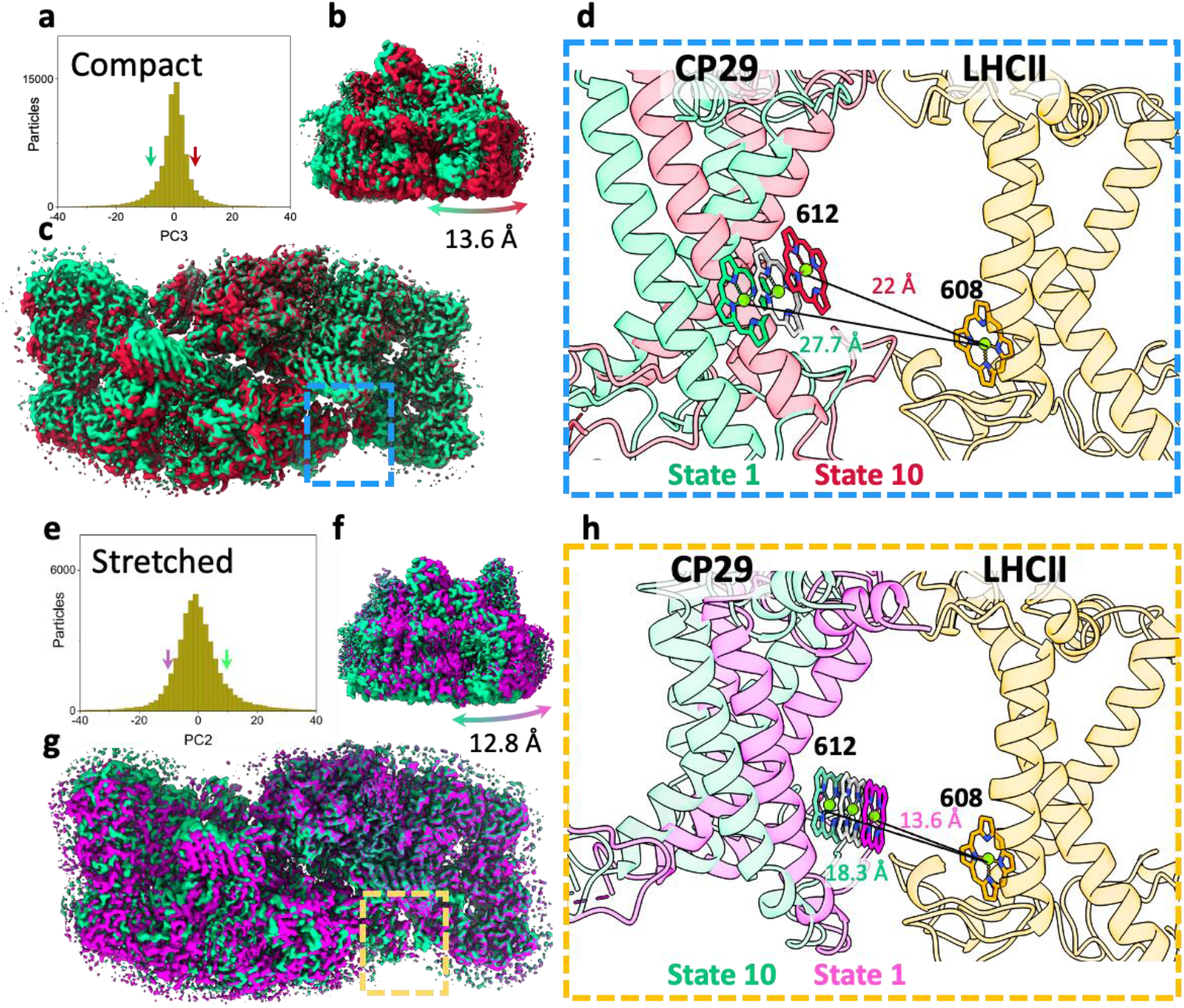
Heterogeneity within PSII states. **a**. Continuous heterogeneity of C2S2_COMP_ particles distribution along the third principal component (PC) axis. Each PC was divided into ten states separated by nine percent of the particle population along the PC axis. State 1 is marked with a green arrow and state 10 with a red arrow. State 1 and 10 are coloured in green and red in panels a-d. **b**. To visualize the state differences, one PSII monomer was superposed, and the other monomer was used to visualize the differences. Membrane plane view of the shift in position of C2S2_COMP_ in the third PC. **c**. Luminal view of the shift in position of C2S2_COMP_. CP29 and LHCII are marked with a blue rectangle **d**. Zoom-in on the change in CP29 position between states 1 and 10. The change in distance between CP29 Chl 612 and LHCII M1 Chl 608 is shown. LHCII M1 (from the superposed PSII monomer) is coloured light orange and the consensus position of Chl 612 is shown in grey. **e**. Continuous heterogeneity of C2S2_STR_ particles distribution in the second PC. State 1 is marked with a magenta arrow and state 10 with a teal arrow. Colours are maintained in panels e-h. **f**. Membrane plane view of the shift in position of C2S2_STR_ in the second PC. **g**. Luminal view of the shift in position of C2S2_STR_. CP29 and LHCII are marked with an orange rectangle. **h**. Zoom-in on the change in CP29 position between states 1 and 10. The change in distance between CP29 Chl 612 and LHCII M1 Chl 608 is shown. LHCII M1 is coloured orange and the consensus position of Chl 612 is shown in grey.

### Water channels and post-translational modifications in *Dunaliella* PSII

More than 2100 water molecules were detected in the C2S2_COMP_ model (**Fig. 3a-b**), the first detailed water molecules structure for a eukaryotic PSII. Overall, water molecules are clearly excluded from the membrane space in the PSII core, however the region occupied by LHC’s show a relatively high number of water molecules in the membrane region. This stems from the presence of several conserved charged amino acids in these antennae and is probably important for the inclusion of such hydrophilic residues within the membrane. We used CAVER^46^ to analyze the structure of internal cavities around the Oxygen Evolving Complex (OEC). As expected from the highly conserved environment around the OEC, the water channels identified previously in the high-resolution cyanobacterial core structure^5,47^ are clearly visible in the eukaryotic PSII, and overlap with the results of the internal cavity analysis, these are shown in **Fig. 3c** and named “Large”, “Narrow” and “Wide”, following^47^. When analyzing the side chains lining the cavities around the OEC, a small hydrophobic patch, highly conserved in prokaryotes and eukaryotes (**Extend Data Fig. 7**), was identified at the beginning of the large channel (**Fig. 3d**). This hydrophobic element may facilitate O_2_ release as part of the catalytic cycle (**Fig. 3d**).

**Fig. 3:**
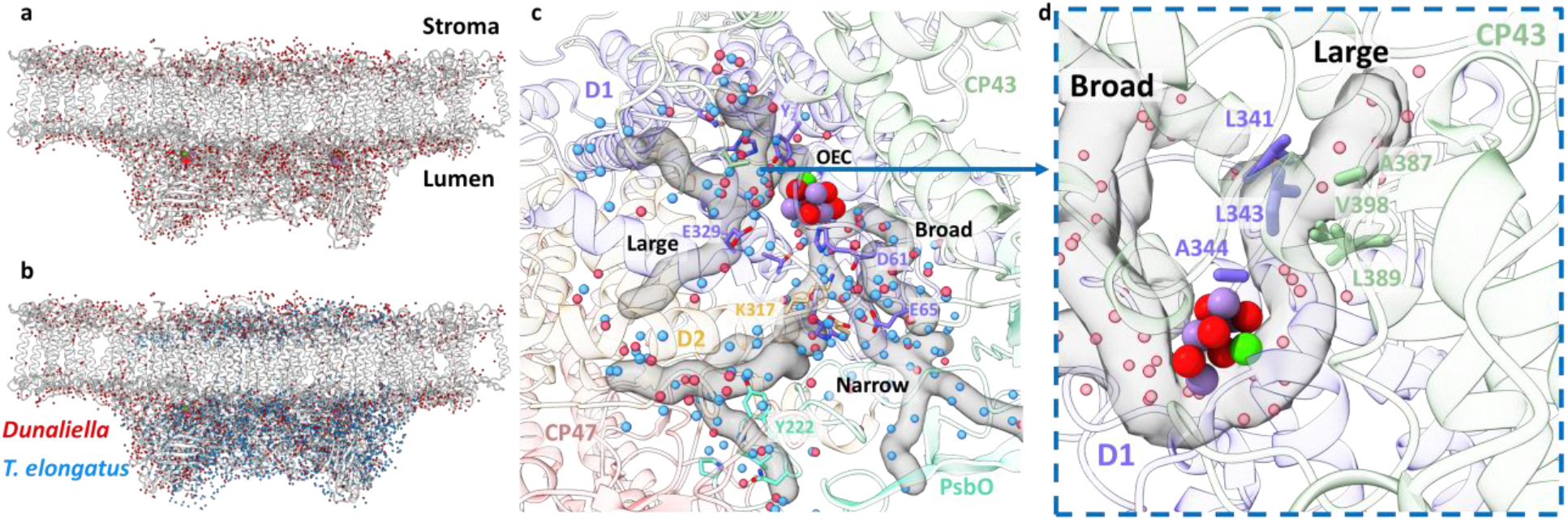
Water distribution and channels in eukaryotic PSII. **a**. Water molecules distribution in *Dunaliella* C2S2_COMP_ structure. The protein scaffold is coloured grey and water molecules are shown as red spheres. **b**. Water molecules distribution in *Dunaliella* C2S2_COMP_ compared to *T. elongatus* PSII core (PDBID 3WU2). *T. elongatus* water molecules are shown as blue spheres. **c**. *Dunaliella* PSII water channels identified by CAVER analysis, shown as grey transparent maps. The Large, Narrow, and Broad channels are annotated along with selected amino acids coloured according to their respective subunits. Water molecules are presented as in panel b. **d**. A hydrophobic patch identified in the large channel near the OEC, which may serve as an O_2_ release pathway. The region shown is indicated by the blue arrow, but the orientation is different to improve visualization.

Several unique map densities were identified during model building, close to the OEC of both configurations a Na^+^ ion was modeled. This Na^+^ ion is coordinated by D1-His337, the backbone carbonyls of D1-Glu333, D1-Arg334, D2-Asn350 and a water molecule, in agreement with the recently identified^48^ binding site (**Extended Data Fig. 8a-b**). This agrees with several studies showing that Na^+^ ions are required for optimal activity of PSII^48,49^. Two additional densities, unique to C2S2_COMP,_ were observed close to CP29-Ser84 and CP47-Cys218. These were modelled as post-translational modifications (PTMs) – Ser84 appears to be phosphorylated and Cys218 seems to be sulfinylated (**Extended Data Fig. 8c-d**). Thus far, PTMs were structurally seen in photosystems only as phosphorylated LHCII bound to PSI during state transition^50,51^. Although they were not identified in-situ, several phosphorylation sites were shown to exist in CP29 large stromal loop^52–55^. CP29 phosphorylation was suggested to be linked with various stress responses, photosynthetic protein degradation and state transition. Cysteine sulfinylation was shown to be linked to superoxide radical (O_2.-_) accumulation, which is subsequently converted by superoxide dismutase (SOD) to hydrogen peroxide (H_2_O_2_) molecules^56–58^. CP47-Cys218 is positioned on the outer edge of PSII, close to the stromal end of the thylakoid membrane, and thus is susceptible to oxidation by H_2_O_2_. The map density around Cys218 suggests two cysteine oxidation events which result in the formation of sulfinic acid (RS-O_2_H).

To summarize, the high-resolution structure of the eukaryotic PSII revealed two distinct states of the PSII complex, adding a new dimension to the known, large compositional heterogeneity of this important system^39,59,60^. The increased map resolution resulted in the identification of PTM’s, and several conserved hydrophobic residues near the OEC, which may serve as a pathway for the release of O_2_. In addition to the two distinct conformations, large levels of continuous structural heterogeneity were discovered within each individual state. Multibody analysis^37^ inherently treats the data as a collection of rigid bodies. This is a good approximation of the heterogeneity in photosynthetic systems but should be regarded as a conservative estimation to additional modes of heterogeneity which exist in this system within each body^42^.

### The structure of *Dunaliella salina* stacked PSII at high resolution

The thylakoid membrane is made of two spatially distinct regions, stroma lamellae and grana stacks, each serving a different role in the photosynthetic process^61,62^. Grana stacks size and numbers are affected by light intensity and ionic composition and can change rapidly^63^. Membrane stacking depends on the presence of cations, mainly Mg^2+^, which is abundant in the thylakoid stroma^64^, and between stacked PSII-LHCII^63^. In vitro, suspending chloroplast membranes in low-salt medium causes grana unstacking, and addition of MgCl_2_ reverts the membranes back to their stacked organisation^65,66^. Several low-resolution cryo-EM models of stacked PSII were obtained in recent years^67–69^, but a high quality PSII structure that can shed light on the contribution of the supercomplex to thylakoid membrane stacking is missing.

The stacked PSII dataset refined to a 3.68 Å map after applying multibody refinement, with each dimer defined as a rigid body. Subsequently, the stacked particles were classified according to the higher quality PSII dimer, and two distinct populations of stacked PSII dimers were obtained, as observed for the unstacked PSII: one in the C2S2_COMP_ conformation solved to 3.36 Å, and the other in the C2S2_STR_ conformation solved to 3.84 Å (**Fig. 4; Extended Data Fig. 9-10**). In both classes the compact conformation exhibited the best fit for the second, lower resolution, PSII dimer.

**Fig. 4:**
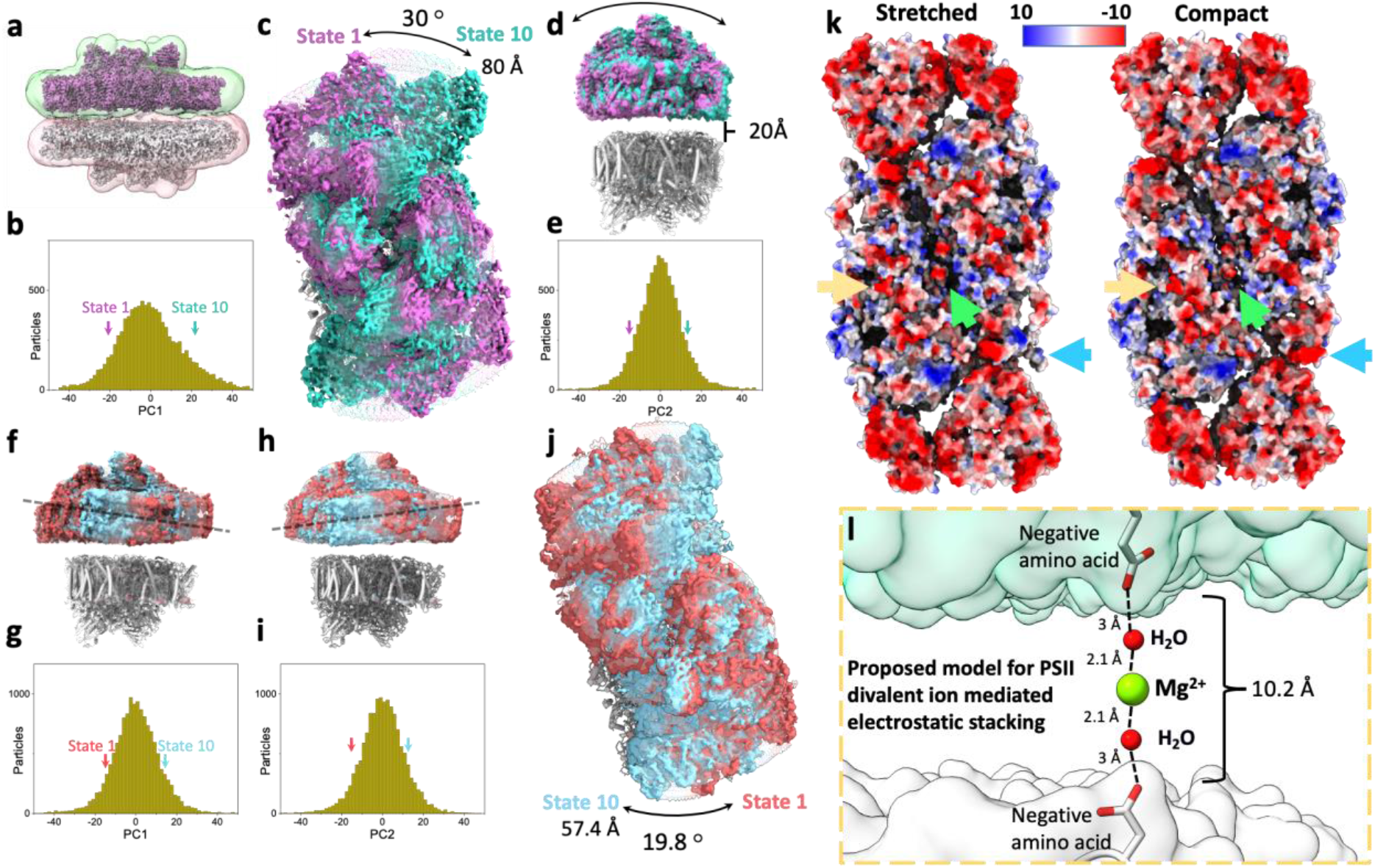
Heterogeneity, electrostatic interactions, and model for PSII stacking. **a**. Stacked *Dunaliella* PSII C2S2_COMP_ maps, and the masks used for multibody refinement. Maps are coloured magenta and grey, and masks in green and red. **b**. The particle distribution along the first PC shows continuous heterogeneity in the stacked C2S2_COMP_. State 1 is marked with a magenta arrow and state 10 with a teal arrow (colours are preserved in panels b-e). **c**. Luminal view of the rotation of the upper PSII dimer between state 1 and state 10 (the bottom dimer was kept in a fixed position). **d**. Membrane plane view of the shift in position of the upper PSII dimer in C2S2_COMP_ second PC. The distance between the upper and lower PSII is shown. **e**. Particle distribution along the second PC of the stacked C2S2_COMP_ shows continuous heterogeneity. **f**. Membrane plane view of the tilt in the upper PSII dimer in C2S2_STR_ particle set first PC. The direction of the tilt is marked with a dashed line. State 1 is coloured red and state 10 in cyan (colours are preserved in panels f-j). **g**. Continuous heterogeneity in the stacked C2S2_STR_ particles distribution along the first PC. **h**. Membrane plane view of the tilt in orientation of the upper PSII dimer in C2S2_STR_ second PC (with the bottom dimer kept fixed). The dashed line shows the tilt axis is opposite to that shown in panel f. **i**. Continuous heterogeneity of stacked C2S2_STR_ particles distribution in the second PC. **j**. Luminal view of the rotation of the upper PSII dimer between state 1 and state 10. **k**. Coulombic electrostatic potential of the stromal region of the stretched (left) and compact (right) conformations. Differences are marked for CP47 C-terminus (orange arrow), the intermonomer space (green arrow) and CP29 (blue arrow). The negative potentials (0 *k*_*B*_T/e > Φ > -10 *k*_*B*_T/e) are coloured red and positive potentials (0 *k*_*B*_T/e < Φ < 10 *k*_*B*_T/e) are coloured blue. **l**. Proposed model for PSII stacking mediated by negatively charged amino acids and Mg^2+^ ions. Upper PSII shown as a green surface and the lower PSII as a white surface.

Roughly 20 Å separate the two stacked dimers in both classes, as previously shown^69^. In several regions this value decreases to approximately 10 Å (**Fig. 4d** and **l**), owing to PSII stromal loops in core subunits and LHCs protruding into the space between the two dimers. Both PSII dimers are shifted by approximately 20 Å relative to each other rather than being perfectly aligned (**Fig. 4a**; **Extended Data Fig. 11a)**. PC analysis showed extensive displacements and rotations across the population with stacked PSII dimers rotating relatively to their opposite dimer by as much as 30°, and shifting by 80 Å in C2S2_COMP_, while in C2S2_STR_ the rotation is more restricted, showing a maximum of 19.8° and a shift of 57 Å (**Fig. 4**). The rotation axis of C2S2_COMP_ appears to be broad region containing the N-termini stromal loops of D2 and CP29 on one dimer, and the stromal loop connecting D2 helices IV and V, CP43 N-terminus, and the C-termini of CP43, CP47 and PsbI on the opposite dimer (**Extended Data Fig. 11**). In the stacked C2S2_STR_ these stacking interactions also include a stromal loop from D1 which is pushed in the stromal gap by a change in the position of the PsbT C-terminus (green arrow in **Fig. 4**), this shift pushes the D1 stromal loop (connecting helices IV and V) into the stromal space and closer to the adjacent dimer (**Extended Data Fig. 11c**). On the axis of rotation which consists of PSII core subunits, additional interactions between different LHCs seem to be essential to maintain stacking. All the rotation states include some degree of LHCs interacting across the stromal gap between opposite PSII dimers and these seem to limit the extent of possible rotational states. In the stacked C2S2_COMP_ particle set the larger range of rotations means that at the extreme states CP26 and LHCII M2 are not involved in stacking interaction and can pair with additional complexes (**Extended Data Fig. 12a**), while in the stacked C2S2_STR_ particle set the smaller rotational range seem to be restricted by CP26 and LHCII M2 interactions (**Extended Data Fig. 12b**). These differences, when repeated over many stacked complexes (with additional LHCII complexes) can translate into substantial changes in thylakoid membrane stacking^70^.

The closest contacts are found at the interface between core subunits from both PSII dimers and CP29, supported by peripheral interactions between LHCII trimer and CP26. Most of the PSII stromal surface is electronegative, and accordingly, most of the amino acids that seem to be involved in stacking interactions are either negatively charged or uncharged (**Fig. 4k**). Interactions spanning 10 Å are probably insufficient to maintain PSII in its stacked arrangement, however if mediated by a Mg^2+^ ion and two-to-four H_2_O molecules, stacking can be stabilized (**Fig. 4l**). These interactions comply with the large degree of rotational freedom observed in the stacked dimers and with the strong dependance of stacked dimers in the presence of Mg^2+^ ions and contribute to thylakoid membrane stacking^66^.

### Summary

The structure of PSII from green algae revealed an unexpected level of conformational flexibility in this highly conserved system. The two stable conformations appear to differ in their antennae connectivity and should be considered in PSII modeling attempts. Within each state, the large degree of structural heterogeneity also contributes to EET and may facilitate transitioning between the different states. In the stacked PSII dimer we do not find any evidence for direct protein interaction connecting the two stacked systems, instead, long range electrostatic interaction can provide the sufficient flexibility needed for the dynamic behaviour of thylakoid membranes.

## Methods

### *Dunaliella* PSII sample preparation

After reaching an OD_730_ of 0.4, the culture was harvested by centrifugation at 4,000 g for 10 min and resuspended in in a medium containing 50 mM HEPES pH 7.5, 300 mM sucrose and 5 mM MgCl_2_. The cells were washed ones in the same buffer and suspended in a buffer containing 25 mM MES, pH 6.5, 10 mM CaCl2, 10 mM MgCl2, 1 M betaine, 5 mM EDTA, 12.5% glycerol. Protease-inhibitors cocktail was added to give final concentrations of 1mM PMSF, 1 µM pepstatin, 60 µM bestatin and 1mM benzamidine. The cells were disrupted by an Avestin EmulsiFlex-C3 at 1,500 psi. Unbroken cells and starch granules were removed by centrifugation at 5,000 g for 5min and the membranes in the supernatant were precipitated by centrifugation in Ti70 rotor at 181,000 g for 1 h. The pellet was suspended in a buffer containing 25 mM MES, pH 6.5, 10 mM CaCl_2_, 10 mM MgCl_2_, 1 M betaine, 5 mM EDTA, 12.5% glycerol giving a Chl concentration of 0.4 mg/ml. n-Decyl-***α***-D-Maltopyranoside (α-DM) was added to a final concentration of 1% and following stirring for 30 min at 4°C the insoluble material was removed by centrifugation of 10,000 g for 5 min. Supernatant was concentrated by centrifugation in TI-75 rotor at 377,000 g for 80 minutes. The pellet was suspended in the above buffer containing 0.3% α-DM at Chl concentration of about 1 mg/ml, loaded on sucrose gradients of 10 to 50% in SW-60 rotor and run at 336,000 g for 15 h. **Extended Data Fig. 13a** shows the distribution of green bands in the tubes. The band containing PSII was concentrated by centrifugation at 550,000 g for 2h and the pellet was suspended in a buffer containing 25 mM MES (pH 6.5), 1 mM CaCl2, 5 mM MgCl2 and 0.1% α-DM to give a Chl concentration of 2 mg Chl/ml. SDS-PAGE of the three bands is presented in **Extended Data Fig. 13b**. The final preparation exhibited oxygen evolution activity of 816 µmole O_2_/mg Chl/h under 560 µmole photons m^−2^ s^−1^ illumination **Extended Data Fig. 13c**.

### Cryo-EM data collection and processing

Concentrated PSII solution (3 µl) was applied on glow-discharged holey carbon grids (Cu Quantifoil R1.2/1.3) that were vitrified for cryo-EM structural determination using a Vitrobot FEI (3 s blot at 4°C and 100% humidity). The images were collected using a 300 kV FEI Titan Krios electron microscope, with a slit width of 20 eV on a GIF-Quantum energy filter, at the EMBL cryo facility, Heidelberg, Germany. A Gatan Quantum K3-Summit detector was used in counting mode at a magnification of 130,000 (yielding a pixel size of 0.64 Å), with a total dose of 51.81 e Å^−2^. EPU was used to collect a total of 13,586 images, which were dose-fractionated into 40 video frames, with defocus values of 0.8–1.9 μm at increments of 0.1 μm. The collected micrographs were motion-corrected and dose-weighted using MotionCor2^71^. The contrast transfer function parameters were estimated using CtfFind v.4.1^72^. A total of 401,467 particles were picked using LoG reference-free picking in RELION3.1^73^. The picked particles were processed for reference-free two-dimensional (2D) averaging. After several rounds of 2D classification, which resulted in 253,804 particles, two initial model was generated using RELION3.1^73^, for the unstacked and stacked PSII.

3D classification of the unstacked PSII revealed two organisations of the light-harvesting complexes surrounding the core complex – C2S and C2S2. C2S contained 21,066 particles were resampled at a pixel size of 0.896 Å, pooled together and processed for 3D homogeneous refinement and multibody refinement^37^ using RELION3.1^73^, giving a final resolution of 3.61 Å. The C2S2 configuration was comprised of 75,904 particles with a C2 symmetry, and these were resampled at a pixel size of 0.896 Å, pooled together and processed for 3D homogeneous refinement and postprocessing using RELION3.1^73^, giving a final resolution of 2.82Å. In an attempt to improve the map density of C2S2, mainly in the vicinity of CP29 and LHCII trimer, 3D classification without refinement was performed, and revealed two distinct C2S2 conformations – compact (C2S2_COMP_) and stretched (C2S2_STR_). C2S2_COMP_ was composed of 39,357 particles that undergone symmetry expansion, 3D homogeneous refinement and multibody refinement^37^ in C1 symmetry to give a final resolution of 2.43 Å, and C2S2_STR_ was composed of 23,014 particles that undergone symmetry expansion, 3D homogeneous refinement and multibody refinement^37^ in C1 symmetry to give a final resolution of 2.62 Å.

23,874 particles that were assigned to the stacked PSII arrangement were resampled at a pixel size of 0.96 Å, pooled together and processed for 3D homogeneous refinement and multibody refinement^37^ in C1 symmetry using RELION3.1^73^ and yielded a final resolution of 3.68 Å. Focused refinement on each individual PSII complex yielded similar resolutions before multibody refinement (3.53 Å and 3.58 Å on each complex), showing both positions are occupied roughly by the same number of complexes. Focused classification was carried out on the upper dimer of the stacked PSII particles to determine if the compact and stretched conformations were also present in the stacked PSII arrangement. This analysis showed that the stacked PSII also contained a mixed population of the compact and stretched conformations. The compact set was comprised of 9,567 particles, and these were pooled together and processed for 3D homogeneous refinement followed by multibody refinement^37^ to give a final resolution of 3.36 Å. The stretched set comprised of 14,307 particles, these were pooled together and processed for 3D homogeneous refinement followed by multibody refinement^37^ to give a final resolution of 3.84 Å. Performing focused refinement on the lower PSII dimer of both conformations suggested a conformation mixture as well, but was less conclusive, due to the lower map quality of the lower PSII dimer and both were fitted with the PSII_COMP_ model (using rigid body refinement) which gave the best overall fit to the map. All the reported resolutions were based on a gold-standard refinement, applying the 0.143 criterion on the FSC between the reconstructed half-maps. (**Extended Data Fig. 2**).

### Model building

To generate the C2S2 PSII, the cryo-EM structure of the C2S2 *Chlamydomonas reinhardtii* PSII model PDB 6KAC^31^ was selected. This model was fitted onto the cryo-EM density map using phenix.dock_in_map in the PHENIX suite^74^, and manually rebuilt using Coot^75^. Stereochemical refinement was performed using phenix.real_space_refine in the PHENIX suite^74^. The final model was validated using MolProbity^76^. The refinement statistics are provided in **Extended Data Table 1**. Local resolution was determined using ResMap^77^, and the figures were generated using UCSF Chimera^78^ and UCSF ChimeraX^79^. Representative cryo-EM densities are shown in **Extended Data Fig. 3**.

## Data availability

The atomic coordinates have been deposited in the Protein Data Bank, with accession code 7PI0 (C2S2_COMP_), 7PI5 (C2S2_STR_), 7PNK (C2S), 7PIN (stacked C2S2_COMP_) and 7PIW (stacked C2S2_STR_). The cryo-EM maps have been deposited in the Electron Microscopy Data Bank, with accession codes EMD-13429 (C2S2_COMP_), EMD-13430 (C2S2_STR_), EMD-13548 (C2S), EMD-13444 (stacked C2S2_COMP_) and EMD-13455 (stacked C2S2_STR_).

## Acknowledgements

Dr Yael Levi-Kalisman is gratefully acknowledged and thanked for vitrifying the samples. We also thank the Electron Microscopy Core Facility (EMCF) at the European Molecular Biology Laboratory (EMBL) for their support and Felix Weis for data collection and excellent technical support. Molecular graphics and analyses were performed with UCSF Chimera, developed by the Resource for Biocomputing, Visualization, and Informatics at the University of California, San Francisco, with support from NIH P41-GM103311. Molecular graphics and analyses performed with UCSF ChimeraX, developed by the Resource for Biocomputing, Visualization, and Informatics at the University of California, San Francisco, with support from National Institutes of Health R01-GM129325 and the Office of Cyber Infrastructure and Computational Biology, National Institute of Allergy and Infectious Diseases. This work was supported by The Israel Science Foundation (Grants No. 569/17 and 199/21), and by German-Israeli Foundation for Scientific Research and Development (GIF), Grant no. G-1483-207/2018.

## Author Contributions

I.C., M.F., Y.M., and N.N. performed the research. I.C., Y.M, and N.N. analyzed the data. I.C., Y.M., and N.N. wrote the manuscript. All the authors discussed, commented on, and approved the final manuscript.

## Author Information

Correspondence and requests for materials should be addressed to N.N or Y.M.

## Competing interests

The authors declare that there are no competing interests.

## Notes

### Competing Interest Statement

The authors have declared no competing interest.

