## Supplementary Information for "Structure of *Dunaliella* Photosystem II reveals conformational flexibility of stacked and unstacked supercomplexes"

### Extended Data Figures

**Extended Data Fig. 1. Cryo-EM data collection and processing scheme for unstacked and stacked PSII complexes. a.** Sample micrograph collected for the *Dunaliella* PSII dataset displaying stacked and unstacked PSII particles from multiple views. **b.** 2D classes showing stacked and unstacked PSII complexes. **c.** Unstacked PSII data processing workflow after 2D classification. An Ab-initio model (coloured cyan) was created in RELION, followed by 3D classification and refinement, resulted in two distinct configuration – C2S (coloured brown-red) with a global resolution of 3.61 Å, and C2S2 (coloured purple) with a global resolution of 2.82 Å. Next, 3D classification (without orientation refinement) was applied to the C2S2 set, and the dataset was separated to two conformations – C2S2 compact (coloured blue) with a global resolution of 2.43 Å, and C2S2 stretched (coloured green) with a global resolution of 2.62 Å. Red arrows show the different CP29 conformation. Particle numbers before C2 expansion are listed, see method section for full details. **d.** Stacked PSII data processing workflow after 2D classification. An Ab-initio model (coloured yellow) was created in RELION, followed by 3D classification and refinement, resulting in a stacked PSII (coloured grey and red) with a global resolution of 3.68 Å. Next, focused 3D classification was applied on the stacked PSII, and the dataset was separated to two conformations according to the upper PSII dimer – stacked C2S2 compact (coloured chocolate and yellow-green) with a global resolution of 3.36 Å, and stacked C2S2 stretched (coloured wheat and sky-blue) with a global resolution of 3.84 Å. Red arrows show the difference between the compact and stretched conformations.

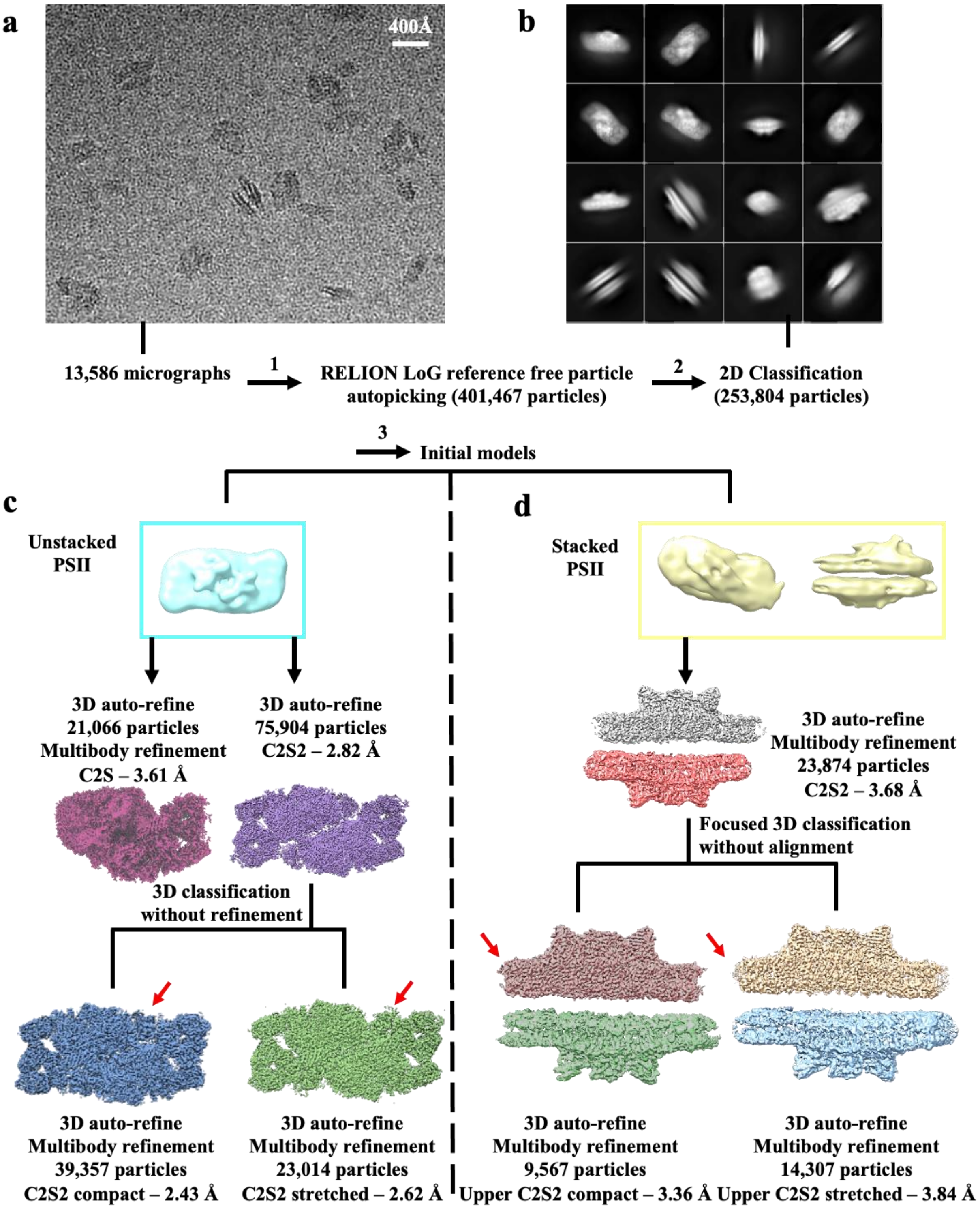
a
400Å
b
13,586 micrographs
1
RELION LoG reference free particle autopicking (401,467 particles)
2
2D Classification (253,804 particles)
3
Initial models
c
Unstacked PSII
d
Stacked PSII
3D auto-refine
21,066 particles
Multibody refinement
C2S – 3.61 Å
3D auto-refine
75,904 particles
C2S2 – 2.82 Å
3D auto-refine
Multibody refinement
23,874 particles
C2S2 – 3.68 Å
Focused 3D classification without alignment
3D classification without refinement
3D auto-refine
Multibody refinement
39,357 particles
C2S2 compact – 2.43 Å
3D auto-refine
Multibody refinement
23,014 particles
C2S2 stretched – 2.62 Å
3D auto-refine
Multibody refinement
9,567 particles
Upper C2S2 compact – 3.36 Å
3D auto-refine
Multibody refinement
14,307 particles
Upper C2S2 stretched – 3.84 Å

**Extended Data Fig. 2. Local resolution and Fourier Shell Correlation (FSC) of *Dunaliella* PSII supercomplexes.** C2S2 local resolution (**a.**) and FSC (**b.**). C2S2 compact local resolution (**c.**) and FSC (**d.**). C2S2 stretched local resolution (**e.**) and FSC (**f.**). C2S local resolution (**g.**) and FSC (**h.**). Stacked C2S2 compact local resolution (**i.**) and FSC (**j.**). C2S2 local resolution (**a.**) and FSC (**b.**). Stacked C2S2 stretched local resolution (**k.**) and FSC (**l.**).

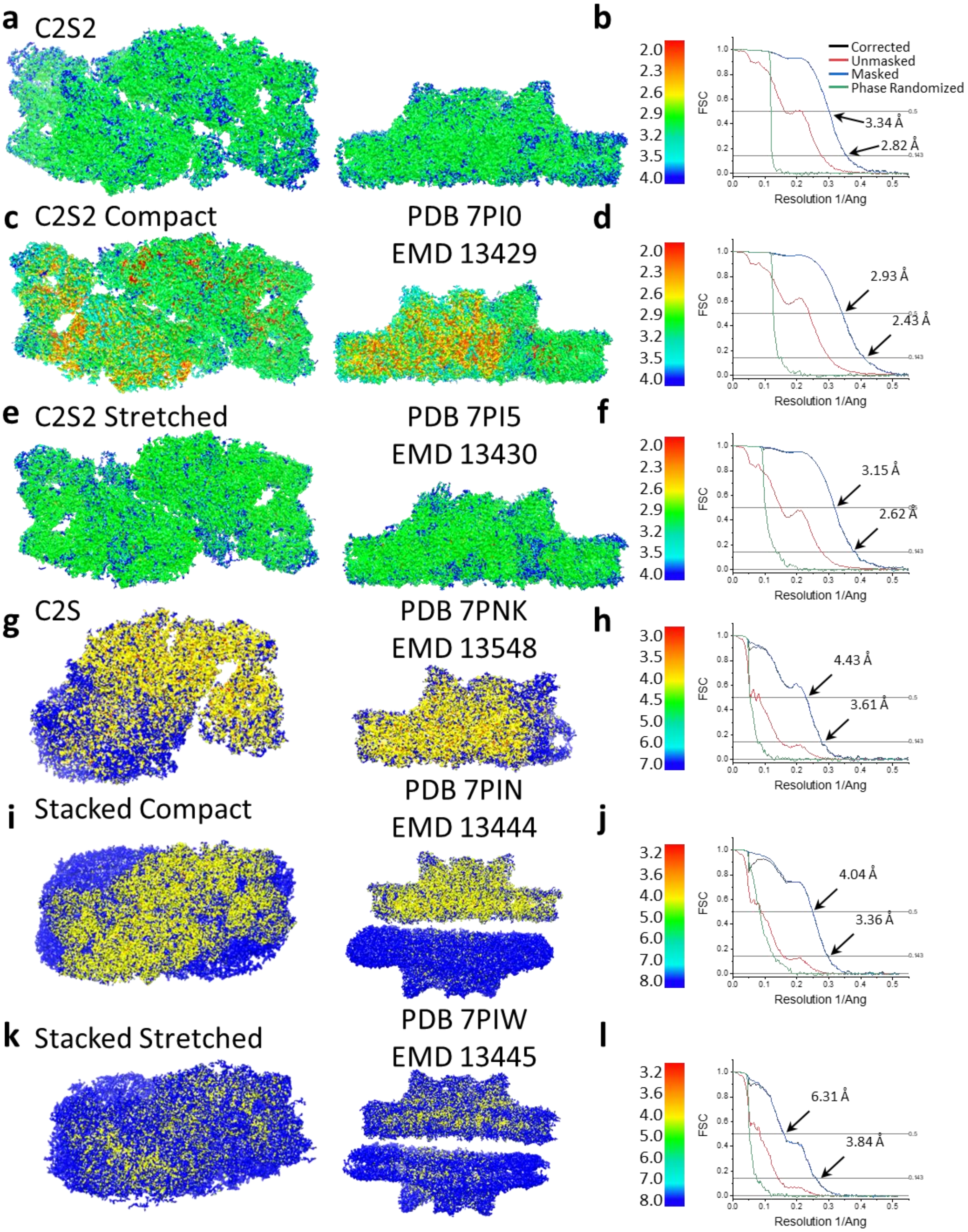

**Extended Data Fig. 3. Map densities of unstacked *Dunaliella* PSII. a.** PSII reaction centre chlorophylls, pheophytins and quinone. Cryo-EM map shown in panels a-h at are contoured at 4σ unless mentioned otherwise. **b.** Cytochrome b559 heme, alpha, and beta subunits. **c.** LHCII M2 Chl b 601. Map shown in 3.5σ contour. **d.** LHCII M2 Chls 602 and 610 alongside coordinating Glu82 and Glu200, Tyr176, Leu83 and Ile201. Water molecules located between a charged and hydrophobic amino acid are shown as red spheres. **e.** PsbH Monogalactosyldiacylglycerol 102. Map shown in 3.5σ contour. **f.** D2 1,2-Dipalmitoyl-Phosphatidyl-Glycerole 409. **g.** CP43 β-carotene 517. **h.** CP26 lutein 621.

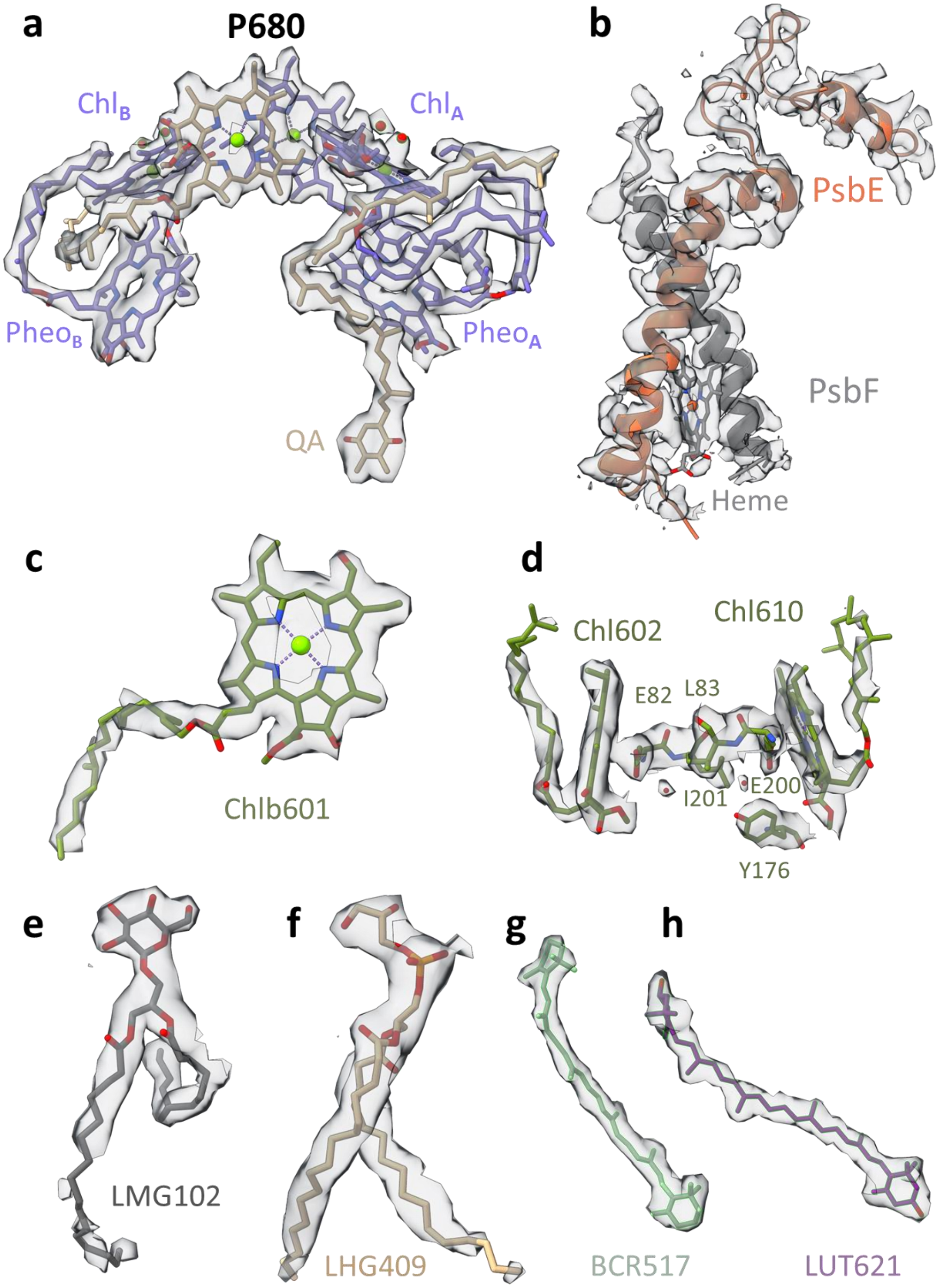

**Extended Data Fig. 4. Changes in CP29 chlorophyll positions between $C2S2_{COMP}$ and $C2S2_{STR}$.** Several CP29 Chl molecules in $C2S2_{STR}$ are positioned close to LHCII and can mediate faster EET from LHCII to PSII, whereas in $C2S2_{COMP}$ CP29 is positioned closer to CP47 and moves away from LHCII. In the $C2S2_{STR}$ conformation, CP29 stromal Chl 612 and Chl 611 are located 16 and 22 Å, respectively, from LHCII M1 Chl 608, while CP29 luminal Chl 613 is located 20 Å from LHCII M1 Chl 605, and CP29 Chl 604 is positioned 23 Å from both LHCII M1 Chl 604 and Chl b 605 (**Fig. 1e**). On the opposite end of CP29, stromal Chl 609 and Chl 603 are found 18 and 22 Å, respectively, from CP47 Chl 616, while luminal Chl b 607 is found 19 Å from CP47 Chl 607. $C2S2_{COMP}$ presents an opposite connectivity – CP29 Chl 612 is located 24 Å from LHCII M1 Chl 608, and CP29 Chl 604 is located 19 and 23 Å from LHCII M1 Chl b 605 and Chl 604, respectively. On the other hand, the rotation of CP29 into the monomer-monomer interface in $C2S2_{COMP}$ set CP29 Chl 603 and Chl 609 19 and 16 Å, respectively, from CP47 Chl 616, while CP29 Chl b 607 is located 15 Å from CP47 Chl 607 (**Fig. 1e**). Chls are presented as spheres, coloured according to their subunit, and the OEC is shown a red and purple sphere.

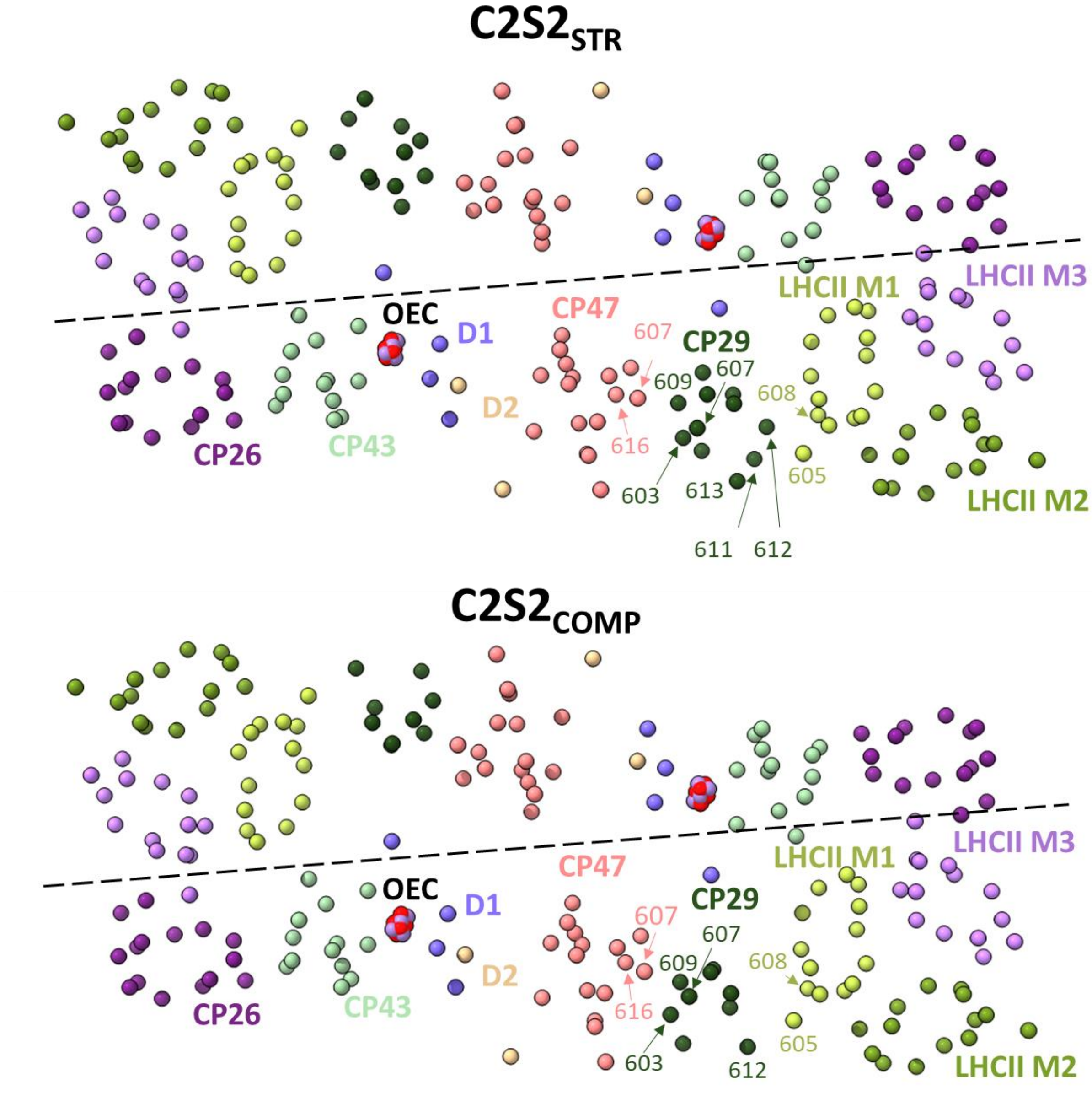

**Extended Data Fig. 5. Principal Component (PC) analysis of C2S2$_{COMP}$.** **a.** Percent of the variance explained by each of the 12 PCs. **b.** The particle distribution along the first six PCs shows continuous heterogeneity in the C2S2$_{COMP}$. States 1 and 10 are marked with a red and green arrow, respectively. **c.** To visualize the state differences, one PSII monomer was superposed, and the other monomer was used to visualize the differences. Luminal view (top), membrane plane view along the long axis (middle), and short axis (bottom) show the shift in position of C2S2$_{COMP}$ in the first PC. **d.** Luminal view (top), membrane plane view along the long axis (middle), and short axis (bottom) show the shift in position of C2S2$_{COMP}$ in the second PC. **e.** Luminal view (top), membrane plane view along the long axis (middle), and short axis (bottom) show the shift in position of C2S2$_{COMP}$ in the third PC.

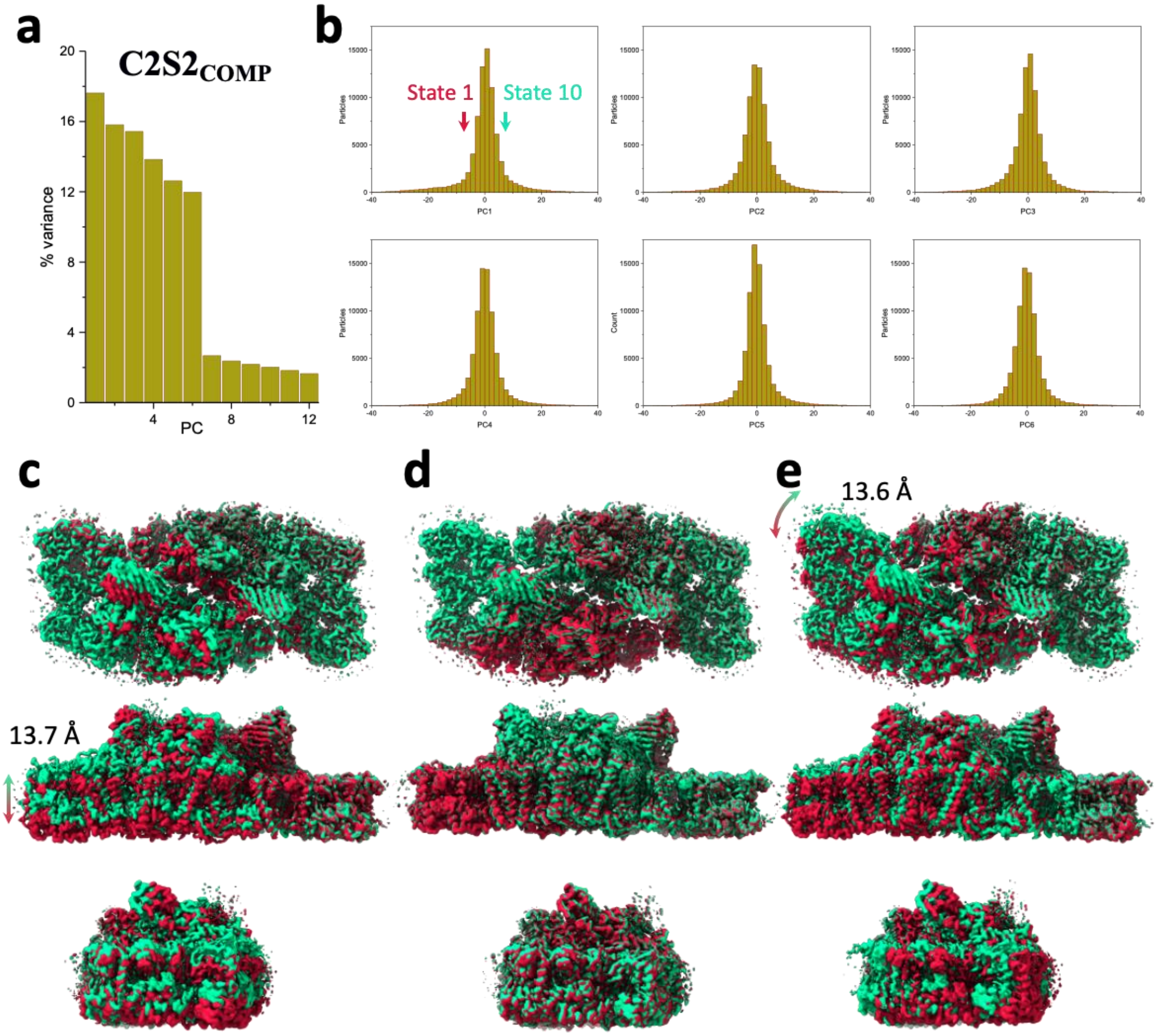

**Extended Data Fig. 6. Principal Component (PC) analysis of $C2S2_{STR}$.** **a.** Percent of the variance explained by each of the 12 PCs. **b.** The particle distribution along the first six PCs shows continuous heterogeneity in the $C2S2_{STR}$. States 1 and 10 are marked with a magenta and teal arrow, respectively. **c.** Luminal view (top), membrane plane view along the long axis (middle), and short axis (bottom) show the shift in position of $C2S2_{STR}$ in the first PC. **d.** Luminal view (top), membrane plane view along the long axis (middle), and short axis (bottom) show the shift in position of $C2S2_{STR}$ in the second PC. **e.** Luminal view (top), membrane plane view along the long axis (middle), and short axis (bottom) show the shift in position of $C2S2_{STR}$ in the third PC.

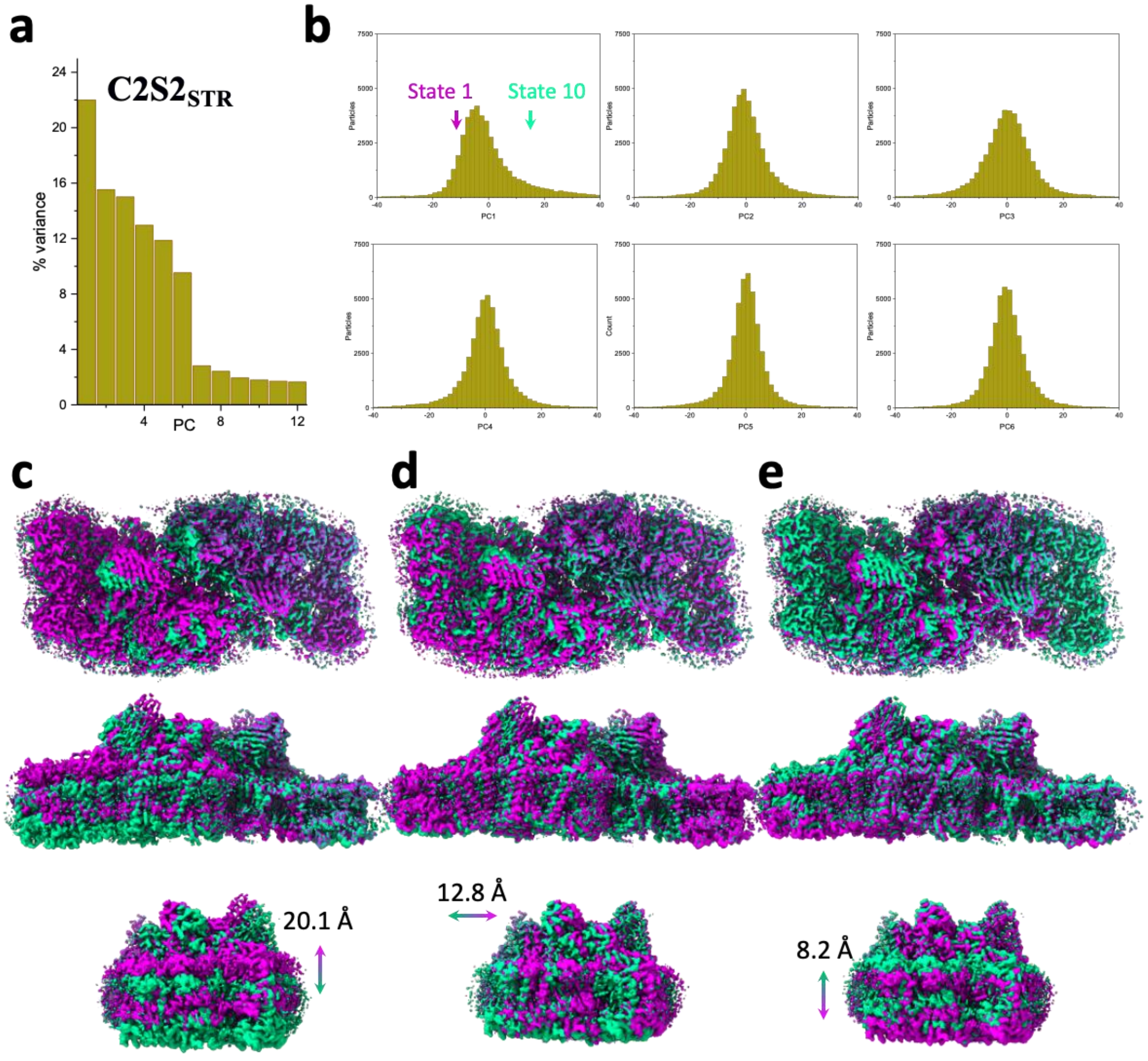

**Extended Data Fig. 7. Sequence alignment of the hydrophobic residues lining the large channel cavity near OEC. a.** Alignment of D1 C-terminus domain, residues 323-344. The hydrophobic residues are marked with black rectangles, together with their conservation score. The organisms used for alignment in panels a and b are *Synechocystis sp. PCC 6803*, *Thermosynechococcus elongatus*, *Nostoc sp. PCC 7120*, *Prochlorococcus marinus*, *Cyanidium caldarium*, *Cyanidischyzon merolae*, *Chaetoceros gracilis*, *Dunaliella salina*, *Chlamydomonas reinhardtii*, *Haematococcus lacustris*, *Pisum sativum*, *Zea Mays and Spinacia oleracea*. **b.** Alignment of CP43 residues 380-408. The hydrophobic residues are marked with black rectangles, together with their conservation score.

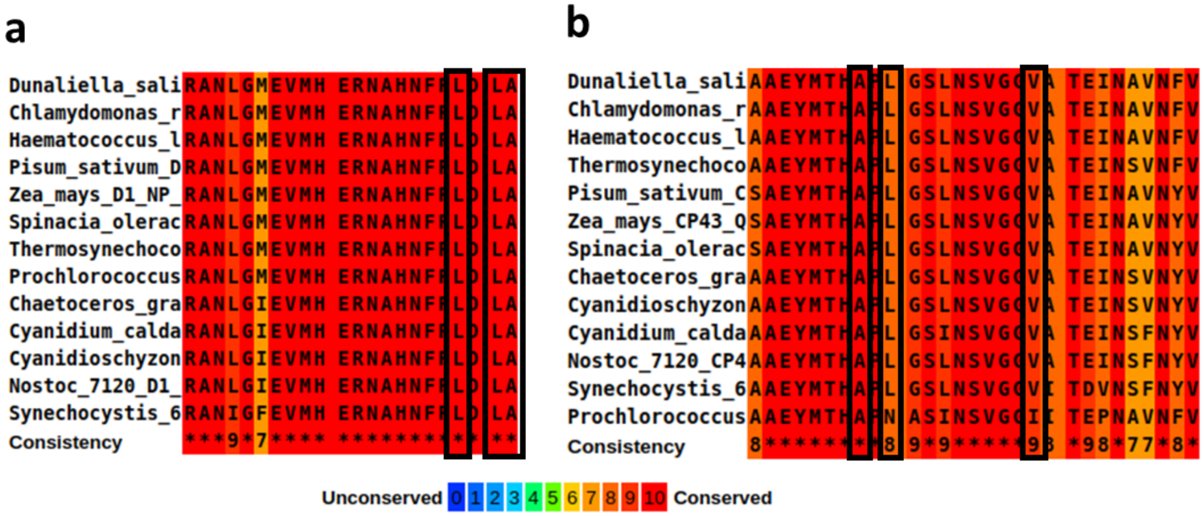

**Extended Data Fig. 8. $Na^+$ ion and PTMs in *Dunaliella* PSII. a.** $Na^+$ binding site close to the OEC. The $Na^+$ ion is coordinated by D1 histidine 337, the backbone carbonyls of D1 glutamic acid 333, arginine 334, D2 asparagine 350 and a water molecule. Amino acids are coloured according to their respective subunits, water molecule as a turquoise sphere, and the OEC coloured red, purple, and green. **b.** Rotated view of $Na^+$ binding site. **c.** CP29 phosphorylated serine 84 and adjacent subunits at the stromal interface. Map surrounding the phosphorylated serine is shown at 3.5σ contour. PsbH lysine 39 coordinating the phosphate is coloured grey, CP29 N-terminal region in dark-green and CP47 in coral. **d.** CP47 sulfinylated cysteine 218 at the stromal region, along with map density at 4σ contour. PsbH histidine 77 coloured grey and CP47 in coral.

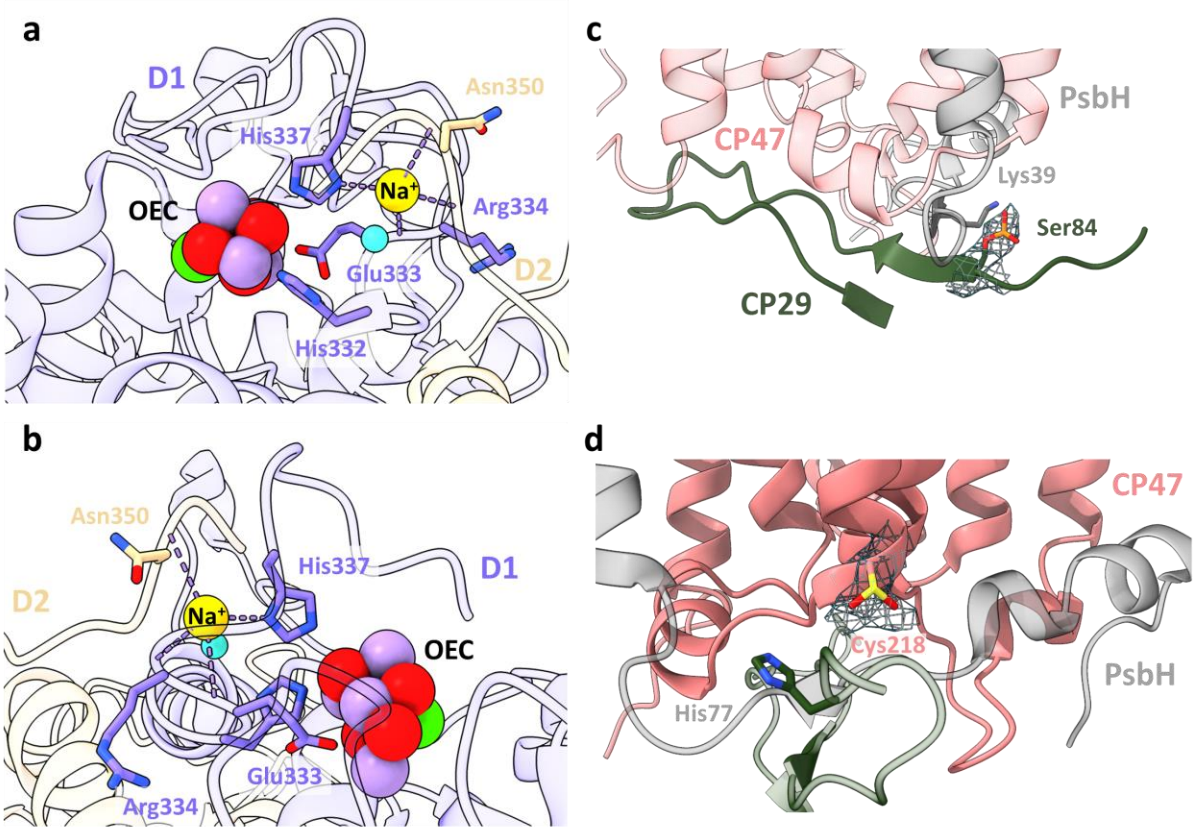

**Extended Data Fig. 9. Principal Component (PC) analysis of stacked C2S2$_{COMP}$. a.** Percent of the variance explained by each of the 12 PCs. **b.** The particle distribution along the first six PCs shows continuous heterogeneity in the C2S2$_{COMP}$. States 1 and 10 are marked with a magenta and teal arrow, respectively. **c.** Membrane plane view along the long axis (top), the short axis (middle), and luminal view (bottom) show the shift in position of stacked C2S2$_{COMP}$ in the first PC. **d.** Membrane plane view along the long axis (top), the short axis (middle), and luminal view (bottom) show the shift in position of stacked C2S2$_{COMP}$ in the second PC. **e.** Membrane plane view along the long axis (top), the short axis (middle), and luminal view (bottom) show the shift in position of stacked C2S2$_{COMP}$ in the third PC.

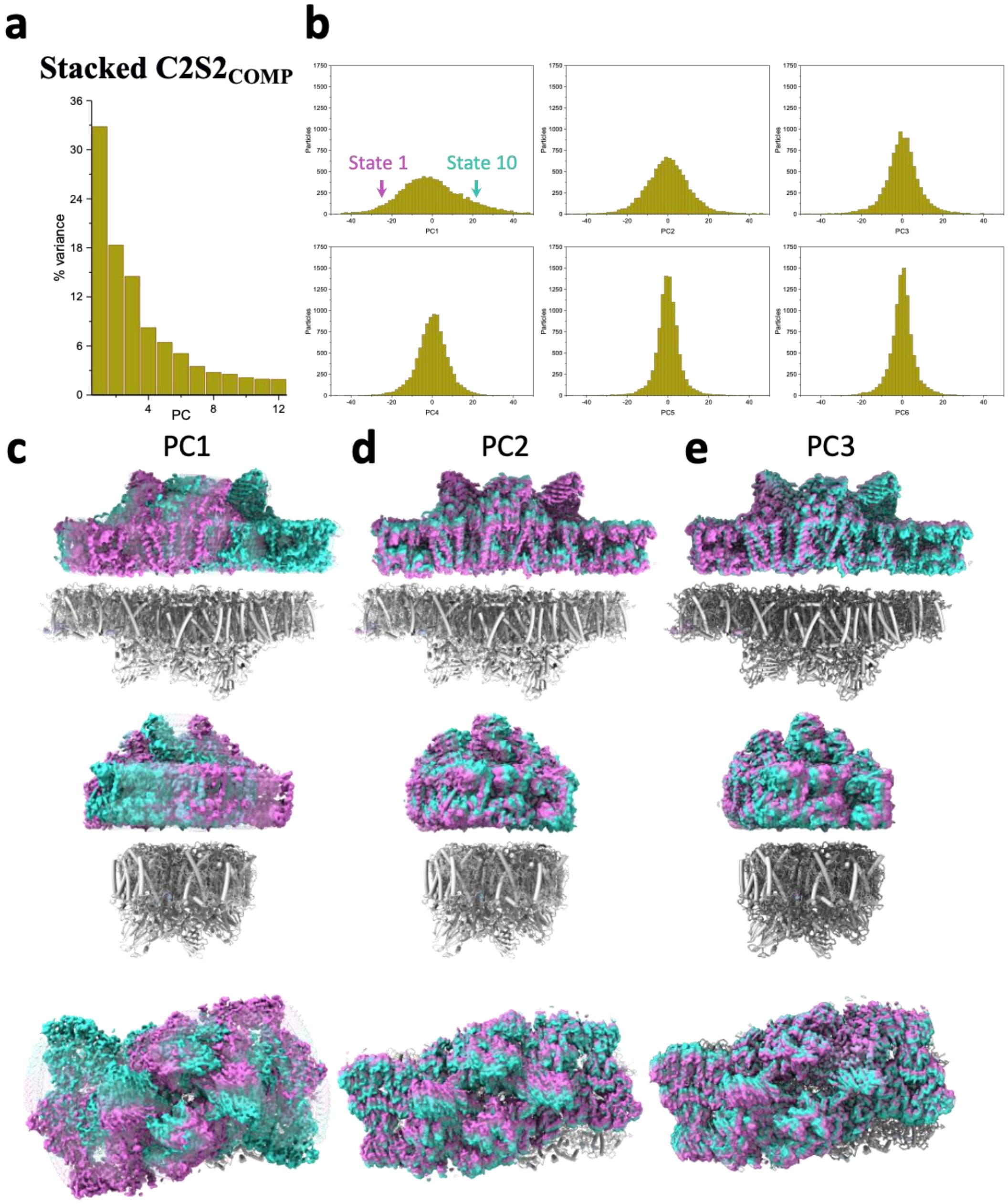

**Extended Data Fig. 10. Principal Component (PC) analysis of stacked $C2S2_{STR}$.** **a.** Percent of the variance explained by each of the 12 PCs. **b.** The particle distribution along the first six PCs shows continuous heterogeneity in the $C2S2_{STR}$. States 1 and 10 are marked with a red and cyan arrow, respectively. **c.** Membrane plane view along the long axis (top), the short axis (middle), and luminal view (bottom) show the shift in position of stacked $C2S2_{STR}$ in the first PC. **d.** Membrane plane view along the long axis (top), the short axis (middle), and luminal view (bottom) show the shift in position of stacked $C2S2_{STR}$ in the second PC. **e.** Membrane plane view along the long axis (top), the short axis (middle), and luminal view (bottom) show the shift in position of stacked $C2S2_{STR}$ in the third PC.

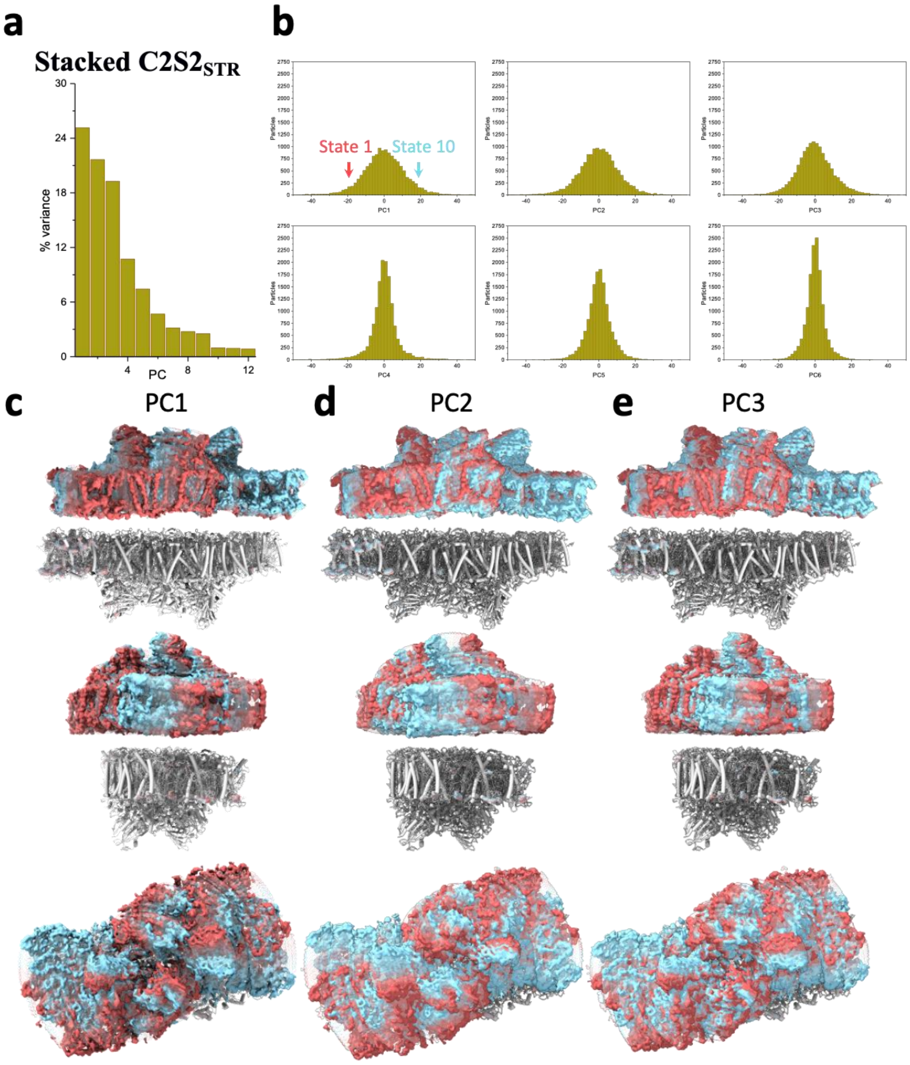

**Extended Data Fig. 11. Stromal interactions between PSII subunits in the stacked configurations. a.** Structure of stacked *Dunaliella* C2S2$_{COMP}$ showing a shift in position of each PSII dimer. The region where PSII core subunits are nearest is marked with a blue rectangle. **b.** Zoom-in on the stromal interactions between PSII$_{COMP}$ dimers, displaying close subunits from both dimers. The map density near CP29 is shown at 2σ, coloured in transparent blue. Subunits are colour coded, and the C and N termini are marked as C and N, respectively. **c.** Zoom-in on the stromal interactions between PSII$_{STR}$ dimers, displaying close subunits from both dimers.

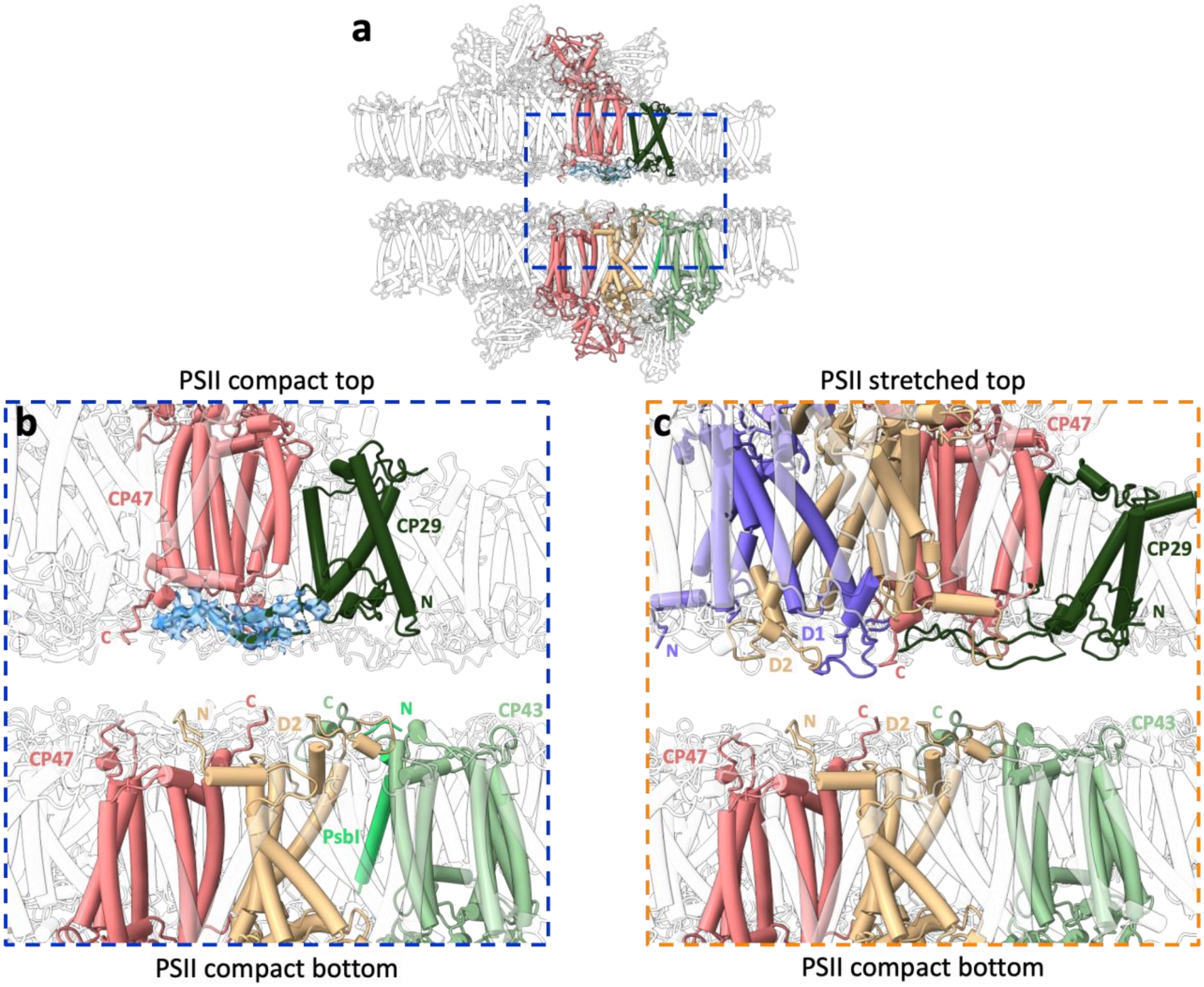

**Extended Data Fig. 12. LHC interactions limit stacked PSII rotation. a**. In stacked C2S2$_{COMP}$ (right), the rotation is confined by LHCII M3 in state 1 (magenta) and 10 (teal). In both conformation the rotation extends from LHCII M2 (state 1) to CP26 (state 10), shown in light grey. **b**. Differences in the extent of PSII rotation are affected by LHC stromal loops. The stacked C2S2$_{STR}$ (left) state 10 (cyan) is tilted compared to the initial position of state 1 (red), bringing the top dimer closer to the lower dimer, resulting in a shorter range of rotations from CP26 in state 1 to LHCII M2 in state 10.

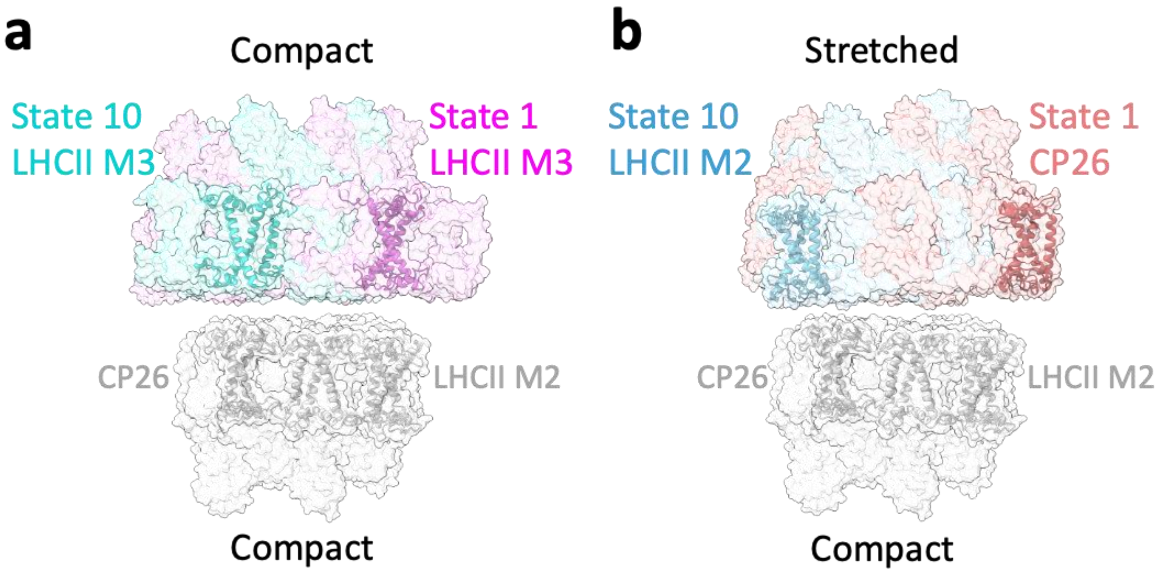

**Extended Data Fig. 13. Sucrose gradient and SDS-PAGE of *Dunaliella* PSII preparation. a.** Sucrose density gradient of the final *Dunaliella* PSII preparation. The three fractions collected are marked As, Bs, and Cs. The Bs fraction was used for cryo-EM data collection. **b.** SDS-PAGE of the three fractions. **c.** Oxygen evolution rates for the collected fractions.

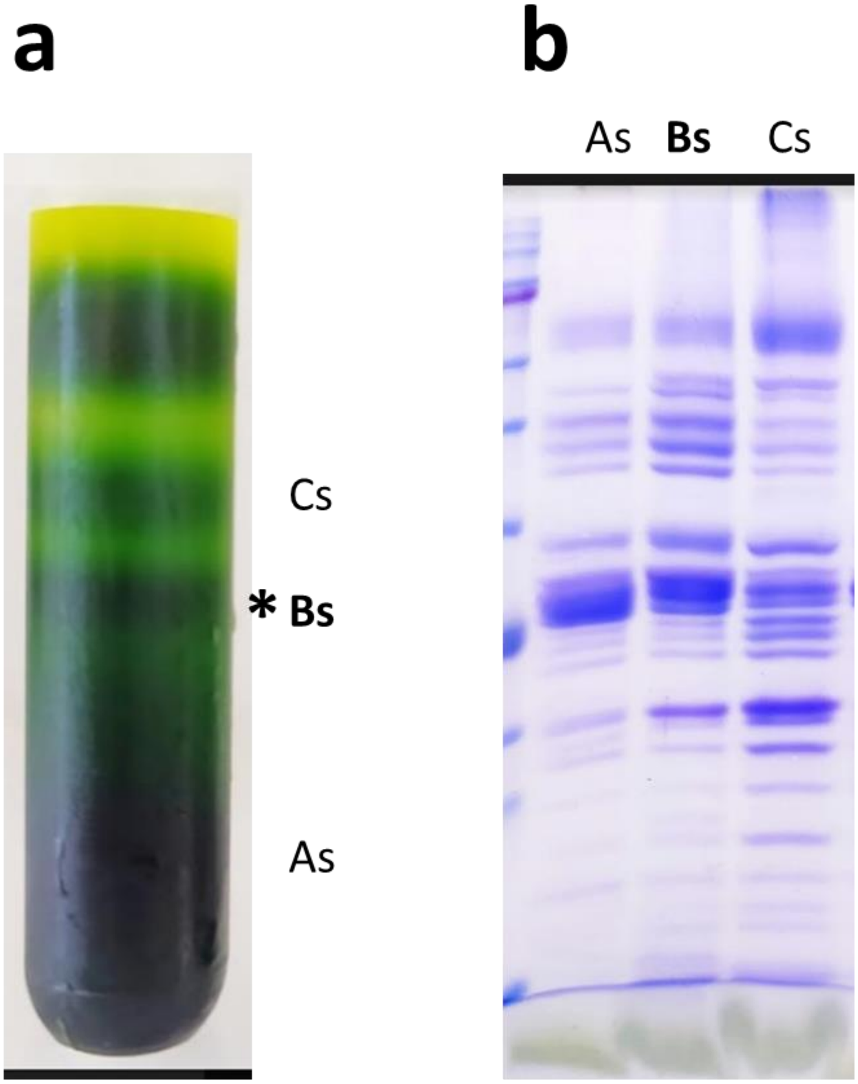

**c**

| Fraction | Oxygen evolution, µmol O2/hour*mg chl |
|---|---|
| As | 276 |
| **Bs** | 816 |
| Cs | 204 |

**Extended Data Table. 1. Cryo-EM data collection, refinement, and validation statistics.**

| | $C2S2_{COMP}$ (EMDB-13429) (PDB 7PI0) | $C2S2_{STR}$ (EMDB-13430) (PDB 7PI5) | C2S (EMDB-13548) (PDB 7PNK) | Stacked $C2S2_{COMP}$ (EMDB-13444) (PDB 7PIN) | Stacked $C2S2_{STR}$ (EMDB-13445) (PDB 7PIW) |
|---|---|---|---|---|---|
| **Data collection and processing** | | | | | |
| Magnification | 130,000 | 130,000 | 130,000 | 130,000 | 130,000 |
| Voltage (kV) | 300 | 300 | 300 | 300 | 300 |
| Electron exposure ($e^{-}$/Å$^2$) | 51.81 | 51.81 | 51.81 | 51.81 | 51.81 |
| Defocus range (μm) | 0.8-1.9 | 0.8-1.9 | 0.8-1.9 | 0.8-1.9 | 0.8-1.9 |
| Pixel size (Å) | 0.64 | 0.64 | 0.64 | 0.64 | 0.64 |
| Symmetry imposed | C1 | C1 | C1 | C1 | C1 |
| Initial particle images (no.) | 401,467 | 401,467 | 401,467 | 401,467 | 401,467 |
| Final particle images (no.) | 39,357 | 23,014 | 21,066 | 9,567 | 14,307 |
| Map resolution (Å) | 2.43 | 2.62 | 3.61 | 3.36 | 3.84 |
| FSC threshold | 0.143 | 0.143 | 0.143 | 0.143 | 0.143 |
| Map resolution range (Å) | 2.0-4.0 | 2.0-4.0 | 3.0-7.0 | 3.2-8.0 | 3.2-8.0 |
| **Refinement** | | | | | |
| Initial model used (PDB code) | 6KAC | 6KAC | 6KAC | 6KAC | 6KAC |
| Model resolution (Å) | 2.70 | 2.70 | 2.70 | 2.70 | 2.70 |
| FSC threshold | 0.143 | 0.143 | 0.143 | 0.143 | 0.143 |
| Model resolution range (Å) | 2.4-5.6 | 2.4-5.6 | 2.4-5.6 | 2.4-5.6 | 2.4-5.6 |
| Map sharpening *B* factor (Å$^2$) | -59.66 | -41.14 | -64.65 | -58.56 | -73.16 |
| Model composition | | | | | |
| Non-hydrogen atoms | 78443 | 76299 | 60673 | 152602 | 151508 |
| Protein residues | 50 | 50 | 44 | 100 | 100 |
| Ligands | 361 | 343 | 250 | 722 | 703 |
| *B* factors (Å$^2$) | | | | | |
| Protein | 14.09-78.39 | 5.21-68.64 | 15.68-156.42 | 16.33-192.00 | 20.51-263.90 |
| Ligand | 17.84-98.34 | 11.99-99.23 | 16.63-92.29 | 20.05-206.46 | 25.08-210.69 |
| R.m.s. deviations | | | | | |
| Bond lengths (Å) | 0.007 | 0.007 | 0.005 | 0.005 | 0.009 |
| Bond angles (°) | 1.662 | 1.596 | 1.513 | 1.632 | 1.681 |
| Validation | | | | | |
| MolProbity score | 1.84 | 1.93 | 1.93 | 1.96 | 2.10 |
| Clashscore | 10.14 | 10.60 | 11.86 | 12.29 | 14.68 |
| Poor rotamers (%) | 0.03 | 0.05 | 0.08 | 0.04 | 0.03 |
| Ramachandran plot | | | | | |
| Favored (%) | 95.48 | 94.35 | 95.06 | 94.73 | 93.53 |
| Allowed (%) | 4.31 | 5.24 | 4.79 | 5.11 | 6.19 |
| Disallowed (%) | 0.20 | 0.41 | 0.15 | 0.16 | 0.27 |

**Extended Data Table. 2. Changes in location of CP29 chlorophylls in the compact and stretched conformations PCs.** Shift in chlorophyll position between state 1 and 10 in the first three PCs of the compact (Comp) and stretched (Str) unstacked PSII. Distances are in Å. The average shift and standard deviation are presented for each component.

| CP29 Chl | 602 | 603 | 604 | 606 | 607 | 608 | 609 | 610 | 612 | Average shift | Stdev |
|---|---|---|---|---|---|---|---|---|---|---|---|
| **Comp PC1** | 4.8 | 3.7 | 2.2 | 2 | 3.3 | 1.3 | 2.4 | 2.2 | 3.5 | **2.8** | **1.1** |
| **Comp PC2** | 0.5 | 0.4 | 0.7 | 0.3 | 0.3 | 0.6 | 0.3 | 0.7 | 0.5 | **0.5** | **0.2** |
| **Comp PC3** | 6 | 4.9 | 5.7 | 5 | 4.6 | 5.4 | 4.6 | 6.4 | 7.3 | **5.5** | **0.9** |
| **Str PC1** | 1.1 | 0.8 | 3.3 | 3.2 | 3 | 2.3 | 0.9 | 2.6 | 2.8 | **2.2** | **1.0** |
| **Str PC2** | 5.7 | 5.3 | 6 | 5.3 | 5.7 | 4.6 | 4.6 | 5.5 | 6.7 | **5.5** | **0.7** |
| **Str PC3** | 3.1 | 2.6 | 3.4 | 3 | 2.9 | 2.9 | 2.5 | 3.5 | 4.1 | **3.1** | **0.5** |

**Extended Data Movie 1. Different conformations of *Dunaliella* C2S2 PSII.** Morph showing the transition from the stretched (magenta) to the compact (green) conformation from a luminal view.

**Extended Data Movie 2. Different conformations of *Dunaliella* C2S2 PSII.** Morph showing the transition from the stretched (magenta) to the compact (green) conformation from a membrane plane view.

**Extended Data Movie 3. Continuous heterogeneity in $C2S2_{COMP}$ PC1.** Transition between all states in $C2S2_{COMP}$ (green) PC1 from a luminal view, showing PSII monomers change in location and orientation along the intermonomer space.

**Extended Data Movie 4. Continuous heterogeneity in $C2S2_{COMP}$ PC3.** Transition between all states in $C2S2_{COMP}$ (green) PC3 from a membrane plane view, showing PSII monomers change in location and orientation along the membrane plane.

**Extended Data Movie 5. Continuous heterogeneity in $C2S2_{STR}$ PC1.** Transition between all states in $C2S2_{STR}$ (magenta) PC1 from a luminal view, showing PSII monomers change in location and orientation along the intermonomer space.

**Extended Data Movie 6. Continuous heterogeneity in $C2S2_{STR}$ PC2.** Transition between all states in $C2S2_{STR}$ (magenta) PC2 from a membrane plane view, showing PSII monomers change in location and orientation along the membrane plane.

**Extended Data Movie 7. Continuous heterogeneity in stacked $C2S2_{COMP}$ PC1.** Transition between all states in stacked $C2S2_{COMP}$ (orchid) PC1 from a luminal view, showing the rotation of the upper PSII dimer compared to the lower dimer.

**Extended Data Movie 8. Continuous heterogeneity in stacked $C2S2_{COMP}$ PC2.** Transition between all states in stacked $C2S2_{COMP}$ (orchid) PC2 from a membrane plane view, showing upper PSII dimer tilting to and from the lower dimer.

**Extended Data Movie 9. Continuous heterogeneity in stacked $C2S2_{STR}$ PC1.** Transition between all states in stacked $C2S2_{STR}$ (red) PC1 from a luminal view, showing the rotation of the upper PSII dimer compared to the lower dimer.

**Extended Data Movie 10. Continuous heterogeneity in stacked $C2S2_{STR}$ PC2.** Transition between all states in stacked $C2S2_{STR}$ (red) PC2 from a membrane plane view, showing upper PSII dimer tilting to and from the lower dimer.
